# A Duplex Structure of SARM1 Octamers Induced by a New Inhibitor

**DOI:** 10.1101/2022.03.02.482641

**Authors:** Tami Khazma, Yarden Golan-Vaishenker, Julia Guez-Haddad, Atira Grossman, Radhika Sain, Alexander Plotnikov, Ran Zalk, Avraham Yaron, Michael Hons, Yarden Opatowsky

## Abstract

In recent years, there has been growing interest in SARM1 as a potential breakthrough drug target for treating various pathologies of axon degeneration. SARM1-mediated axon degeneration relies on its TIR domain NADase activity, but recent structural data suggest that the non-catalytic ARM domain could also serve as a pharmacological site as it has an allosteric inhibitory function. Here, we screened for synthetic small molecules that inhibit SARM1 by stabilizing the ARM-dependent inactive, compact octamer ring conformation, and tested a selected set of these compounds in a DRG axon degeneration assay. Using cryo-EM, we found that one of the newly discovered inhibitors, a Calmidazolium designated TK106, not only stabilizes the previously reported inhibited conformation of the octamer, but also promotes the formation of a meta-stable structure: a duplex of octamers (16 protomers), which we have now determined to 4.0 Å resolution. In the duplex, each ARM domain protomer is not only engaged in lateral interactions with neighboring protomers but is further stabilized by contralateral contacts with the opposing octamer ring. Mutagenesis of the duplex contact sites leads to SARM1 activation in cultured cells. Based on our data we propose that the duplex assembly constitutes an additional auto-inhibition mechanism that tightly prevents pre-mature activation and axon degeneration.

## Introduction

SARM1 induces axonal degeneration after nerve injury (Gerdts et al., 2013; Osterloh et al., 2012) and in other neuropathological conditions, such as chemotherapy induced peripheral neuropathy (CIPN) (Cetinkaya-Fisgin et al., 2020; Geisler et al., 2016). Consistent with its pro-degenerative function, genetic ablation of SARM1 protects against axonal and other types of neuronal degeneration, without inflicting any apparent impediments on the animal models (Kim et al., 2007; Ko et al., 2020; Ozaki et al., 2020; Sasaki et al., 2020; Turkiew et al., 2017; Uccellini et al., 2020). The realization that pharmacological targeting of SARM1 might constitute a new safe and effective therapy against a variety of axonopathies drives a pursuit to identify and develop SARM1 inhibitors (Bosanac et al., 2021; Hughes et al., 2021; Li et al., 2021; Loring et al., 2020). Because SARM1 involvement in axonal degeneration relies entirely on its ability to hydrolyze NAD+ (Essuman et al., 2017), small molecules that directly compromise this enzymatic activity are regarded as SARM1 inhibitors. However, recent structural investigations of the full-length SARM1 have revealed a mechanism of allosteric regulation over the NADase activity of SARM1 that might be exploited in the design of more sophisticated inhibitors (Tong, 2021).

The basic topology of SARM1 includes an N-terminal ARM domain, followed by two SAM and one TIR domain (Figure 1), which mediate auto-inhibition (Chuang and Bargmann, 2005; Summers et al., 2016), oligomerization (Gerdts et al., 2013), and NADase activity (Gerdts et al., 2015), respectively. The NADase activity depends on high proximity between at least two TIR domains, as demonstrated by induced dimerization of TIR, which results in NAD+ hydrolysis (Gerdts et al., 2015; Gerdts et al., 2016). In our first report in which we probed the structure and domain organization of SARM1 (Sporny et al., 2019), we determined the x-ray crystal structure of the tandem SAM domains (as did (Horsefield et al., 2019)). Using negative stain single particle EM to visualize the intact protein, we found that the ARM and TIR domains are closely arranged around an inner ring octamer of the tandem SAM domains. Later, using cryo-EM, we (Sporny et al., 2020) and others (Bratkowski et al., 2020; Figley et al., 2021; Jiang et al., 2020; Li et al., 2021; Shen et al., 2021) determined the high-resolution structure and revealed the molecular details of the ARM-SAM-TIR inter-domains interfaces. The tightly packed structure in all the above reports points at an auto-inhibitory conformation, as the TIR domains are kept separated from each other in a way that presumably prevents NADase activity. Thus, activation of SARM1 probably involves a conformational re-arrangement that would allow nearing of the TIR domains. We further investigated what keeps SARM1 in auto-inhibited conformation in vivo and what triggers domain re-arrangement and NADase activation. We found that NAD+ binds to the concave surface of the ARM domains (as did (Jiang et al., 2020)) and, via an allosteric substrate inhibition mechanism, inhibits SARM1 NADase activity by inducing the compact, auto-inhibited conformation (Figure 1B). Importantly, mutations at the allosteric site that interfere with NAD+ binding, such as W103D (but not W103A), L152A, R157E, R322E (Sporny et al., 2020), or R110E-R157E-K193E (Jiang et al., 2020), elevate hSARM1 activity in cultured cells and axons, demonstrating that the apo form is largely activated, and that an inhibitory compound (such as NAD+) must be included to prevent pre-mature activation. Other reports have shown that different nicotine nucleotide derivatives modulate SARM1 activity, and can bind to an equivalent allosteric site in the Drosophila ARM domain. While nicotinic acid mononucleotide (NaMN), like NAD+, inhibits hSARM1 activity (Sasaki et al., 2021), nicotinamide mononucleotide (NMN) inflicts further activation (Bratkowski et al., 2020; Figley et al., 2021; Liu et al., 2018a; Zhao et al., 2019). Likewise, synthetic molecules where also shown to activate the NADase activity of SARM1 (Loreto et al., 2020; Wu et al., 2021; Zhao et al., 2019). A synergistic model was proposed in which the ratio between the axonal concentrations of NAD+ (and possibly also NaMN) and NMN dictates the balance between inhibition (high NAD+ or NaMN and low NMN) and activation (low NAD+ or NaMN and high NMN) of hSARM1 (Figley et al., 2021; Sasaki et al., 2021) (Figure 1B). However, in spite of the wealth of structural information, it is still a puzzle how the binding of NAD+ and the other nicotine nucleotide derivatives to the allosteric site at the ARM domain inflicts the structural rearrangement thought to control hSARM1 activation. Specifically, the most obvious structural differences within the ARM domain between apo and ligand-bound forms are at the ARM^1^-ARM^2^ loop (res. 310-320) and to a lesser extent, the N’ terminal ARM^1^ **α**1 helix; none of which are directly engaged in domain-domain contacts within the octamer monoplex.

**Figure 1.**
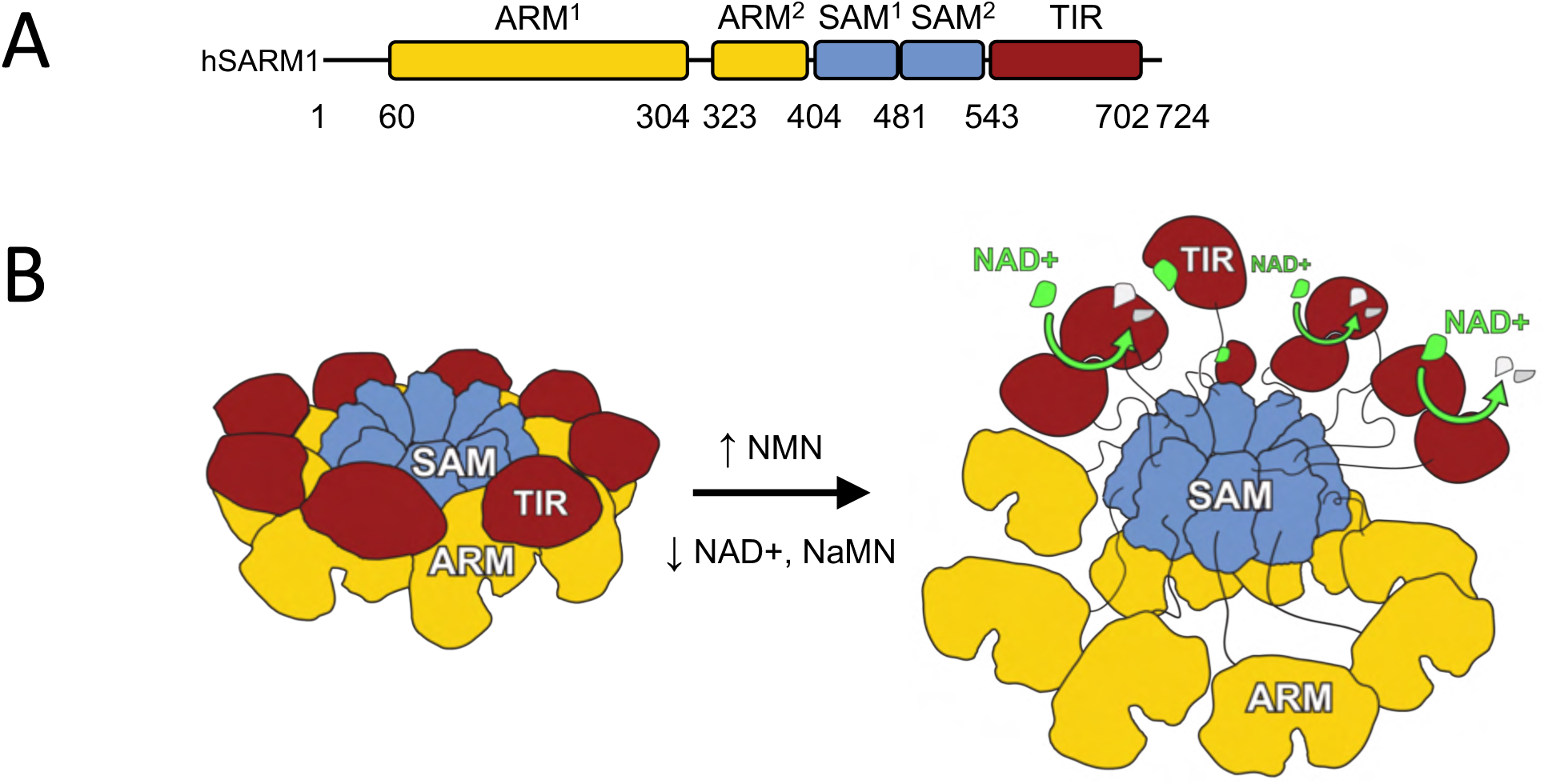
Domain organization, structure, and activation model of hSARM1. A) Color-coded organization, nomenclature, and boundaries of the hSARM1 ARM, SAM, and TIR domains. B) A model for hSARM1 inhibition and activation. Color-coded organization as in (A). A simplified representation of the hSARM1 3D structure octamer is presented on the left. In this structure, which is based on high-resolution cryo-EM reconstructions (Sporny et al., 2020), an inner, stable octamer ring of SAM domains is surrounded by a transient peripheral ring of ARM and TIR domains. TIR domains need to be closely dimerized to gain NADase activity. Because each TIR is docked on two neighboring ARM domains, separated from other TIRs, this structure is thought to represent an auto-inhibited conformation. Thus, SARM1 activation (as in the right panel) is thought to involve a conformational re-arrangement that allows TIR dimerization. NAD+, NMN, and NaMN-all can bind to an allosteric site at the concave surface of the ARM.While NAD+ and NaNM inhibit NADase activity, NMN promotes it.

Here, we resolved the structure of hSARM1 in back-to-back duplex form, a total of 16 protomers, at 4.0 Å resolution that represents a meta-stable conformation of hSARM1. The duplex formation is induced by a new non-competitive Calmidazolium inhibitor that we have identified. The contralateral duplex contacts involve molecular sites that two of them are also affected by ligands binding to the ARM domain: ARM^1^ **α**1 helix, ARM^2^ **α**2-**α**3 loop, and a region proximal to the ARM^1^-ARM^2^ loop (Figure 1 - figure supplement 1). When the contralateral contacts are compromised by site-directed mutagenesis, hSARM1 NADase activity in cultured cells is elevated. We also show that the Calmidazolium compound, like some of the other competitive inhibitors that we have identified using a high throughput hSARM1 inhibitor screen assay, inhibit the NADase activity of hSARM1 *in vitro*, but not that of the zebrafish ortholog, thereby demonstrating pharmacological species specificity. Finally, by implementing a well-established mouse DRG *ex vivo* assay, we show that these inhibitors phenocopy the SARM1 null mutant phenotype by delaying axon degeneration after axotomy.

## Results

### High-throughput screening for SARM1 inhibitors and reciprocal HPLC assay

To discover new and effective inhibitors for hSARM1, we applied a modified fluorescence assay for high-throughput screening. The adaptations introduced in this assay aim to achieve: stable fluorescent signal over time, good sensitivity for inhibition by screened compounds, high signal-to-noise, good reproducibility, accuracy, and diminishing of false-positive readouts. The assay is based on the resazurin in vitro application, as reported in (Sporny et al., 2020), and measures NAD+ hydrolysis by isolated hSARM1. The basic principle of the assay is that the coupled activities of alcohol dehydrogenase and diaphorase, which convert resazurin to the fluorescent resorufin, depend on the amount of available NAD+. Pre-incubation of the NAD+ component with active hSARM1 NADase reduces the amount of NAD+ and, by that, the output fluorescent signal. Introducing a hSARM1 inhibitory compound in the hSARM1-NAD+ incubation stage will result in less hSARM1 NADase activity, more available NAD+ for the alcohol dehydrogenase and diaphorase coupled reaction, and more fluorescence output (Figure 2A). Undesired inhibition of either alcohol dehydrogenase or diaphorase by the screened compounds cannot result in more fluorescence, and therefore, does not produce false-positive signals. To establish a working protocol for high-throughput screening, we looked for optimum hSARM1 and starting NAD+ concentrations that would allow confidently identifying as little as 20% inhibitory activity by the screened compounds. We found that hSARM1 at 12.5nM and NAD+ at a range of 0–250nM give the best performance. In this way, the fluorescence signal trendlines do not overlap at any time-point for over an hour, allowing distinguishing between small differences in NAD+ concentrations (Figure 2B and C). Stopping the further development of the fluorescence reaction by the addition of the dehydrogenase inhibitor iodoacetamide keeps the signal stable for at least another hour (Figure 2D). For control, we used 1mM nicotinamide (NAM), which proved to inhibit SARM1 NADase activity in a dose-response manner (Figure 2E) (Essuman et al., 2017). We note that NAM inhibition of SARM1 was recently put into question (Angeletti et al., 2021), suggesting that instead of preventing NAD+ catalysis it actually participates in a base exchange reaction, but for our practical purpose - to measure the available NAD+ level, it makes no difference. After establishing the working conditions and protocol of the assay, we applied it for high-throughput screening in the Nancy and Stephen Grand Israel National Center for Personalized Medicine. We screened ~150,000 compounds from commercially available libraries of small molecular weight compounds. Out of the 120 molecules that showed at least a 30% reduction in fluorescent signal, 31 molecules were selected, based on pharmacokinetics, drug-likeness and medicinal chemistry properties (Daina et al., 2017). Thirty more molecules that were not picked up by the screening but closely resembled the best hits, were also pursued. Together, these 61 molecules were validated for inhibition with a reciprocal HPLC assay (Figure 3D), which includes 10µM of the inhibitor compound, 50µM of NAD+, and 250nM of hSARM1. Under these conditions, 7 compounds that were also re-validated by mass spectroscopy analysis, inhibited ≥50% of NAD+ hydrolysis activity, compared to control. These 7 compounds were further analyzed for *in vitro* SARM1 inhibition (IC_50_ and Lineweaver Burk plots) and in an *ex vivo* mouse DRG assay for Wallerian degeneration after axotomy.

**Figure 2:**
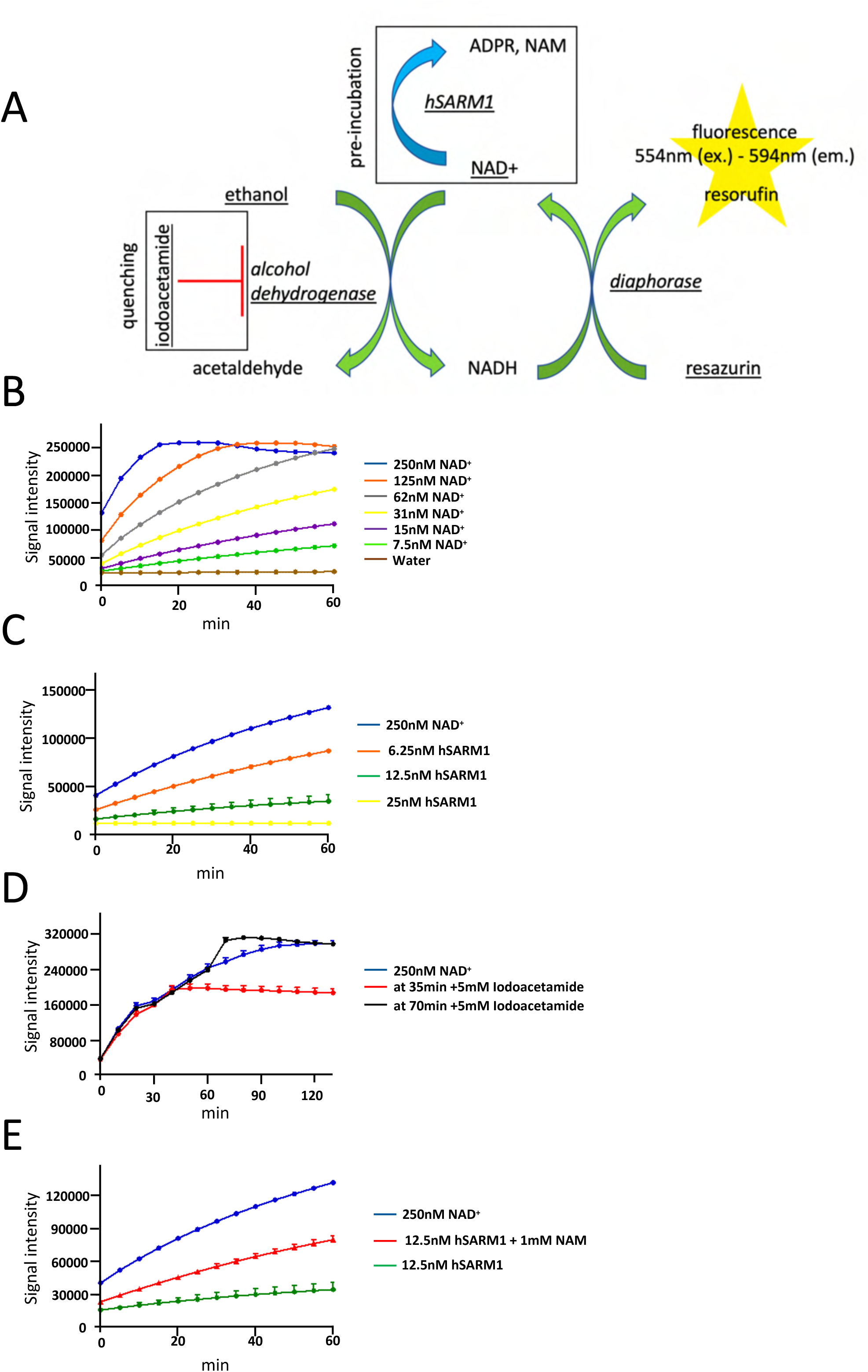
Optimization of resazurin florescence assay for high throughput screening (A) First, NAD+ is pre-incubated with hSARM1 along with each one of the screened compounds. Depending on the NADase activity of hSARM1 in the presence of a particular compound, NAD+ will be consumed, e.g. a complete inhibition of hSARM1 will leave all the NAD+ intact. Next, the underscored components of the enzymatic coupled reaction are added, and eventually, a fluorescent signal is gained. High fluorescence indicates high NAD+ levels and hSARM1 inhibition, and low fluorescence indicates hSARM1 free of inhibition. If reaction quenching is required, the alcohol dehydrogenase inhibitor iodoacetamide will be added. B) Progression of fluorescent signal (measurement every 5 minutes for an hour. Data are presented as mean ± SEM for sixteen repeats) over time in response to various NAD+ inputs. The assay distinguishes between NAD+ levels in the 1-250 nM range up to 25 minutes after reaction commencement. C) Iodoacetamide inhibits alcohol dehydrogenase activity, and so fixes the fluorescence output signal for >100 minutes. Data are presented as mean ± SEM for four repeats D) Pre-incubation of NAD+ with hSARM1 demonstrates a hSARM1 dose response on fluorescence output signal. Data are presented as mean ± SEM for four repeats. E) Fluorescence output responds to hSARM1 inhibition by 1mM of NAM (pre-incubation for 10 minutes at room temperature). Data are presented as mean ± SEM for four repeats.

**Figure 3:**
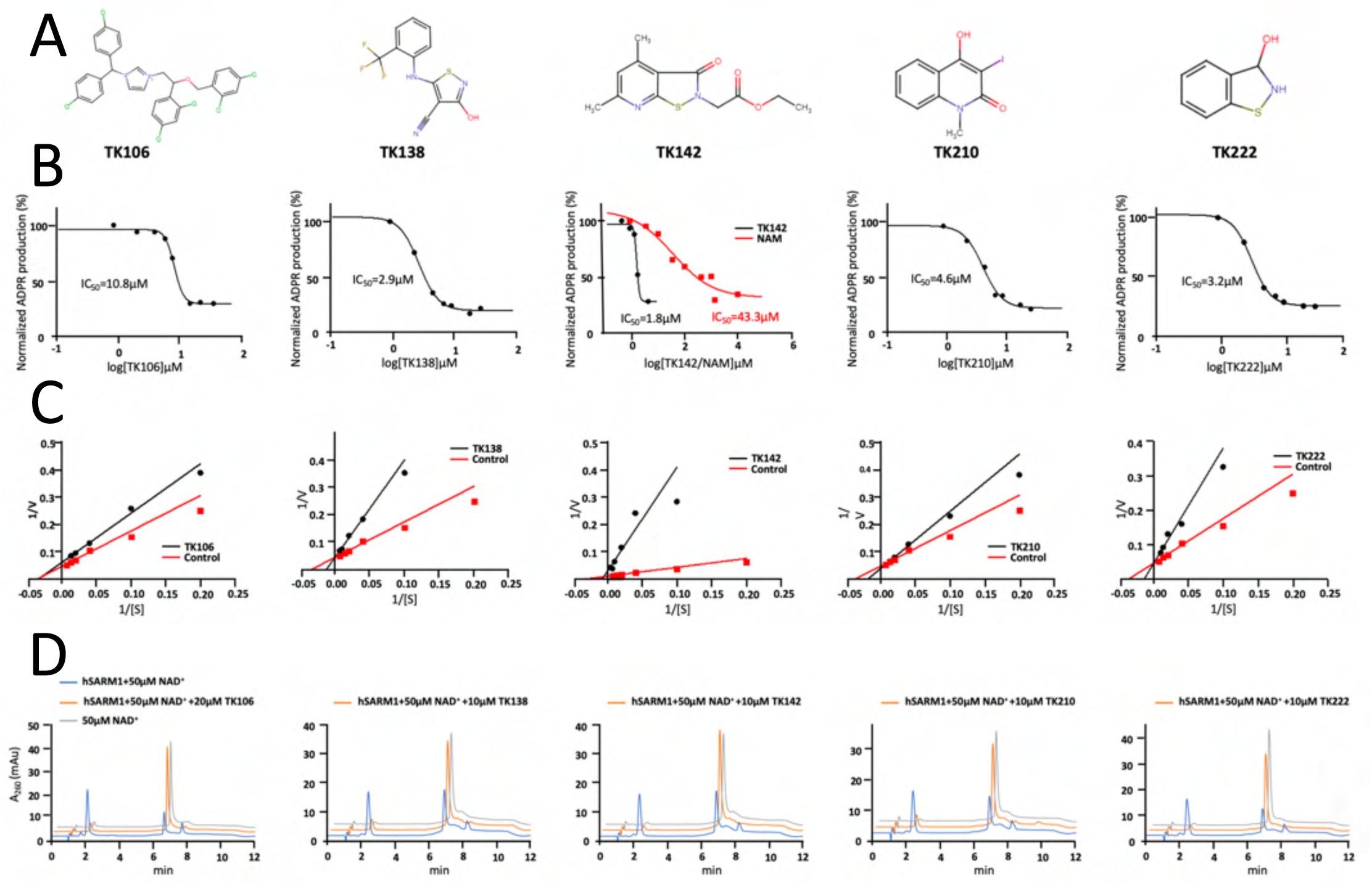
Potency and mode of inhibition of hSARM1 by 5 chemical compounds. A) Chemical structure of hSARM1 inhibitory compounds. B) Determination of IC_50_ values. Various concentrations of the compounds were pre-incubated with hSARM1 for 10 minutes at room temperature, followed by the addition of 50 µM NAD+, initiating the NADase activity. All the assays were carried out at 37 °C for 10 minutes. For reference, the IC_50_ values of the compounds were compared to nicotinamide (NAM), a known hSARM1 inhibitor (see overlaid on the TK142 plot). C) Lineweaver-Burk plots for determination of the mode of inhibition. The NADase activity of hSARM1 was measured at different NAD+ concentrations with and without the presence of 10µM of the five inhibitors. The results show that TK106 inhibits hSARM1 in a non-competitive manner, while the other molecules inhibit hSARM1 in a competitive manner. D) Exemplary HPLC chromatograms. To determine the IC_50_ values, the amount of ADPR production was measured by HPLC and calculated and plotted by GraphPad.

### *in vitro* inhibitors characterization - IC_50_, Lineweaver-Burk plots, and species specificity

At that point, we have focused our analysis on seven SARM1 inhibitory compounds (designated TK###, and presented in SMILES format), the following three of which were picked up in the screening:

TK138) [H]OC1=NSC(N([H])C2=CC=CC=C2C(F)(F)F)=C1C#N

TK142) CCOC(=O)CN1SC2=NC(C)=CC(C)=C2C1=O

TK174) O=C(N1C[C@H]2OC(=O)N(CCCN3CCOCC3)[C@H]2C1)c4ccccn4

And four others were based on similarity to other “hit” compounds that were identified in the screen:

TK106) [Cl-].Clc1ccc(cc1)C(c1ccc(Cl)cc1)n1cc[n+](CC(OCc2ccc(Cl)cc2Cl)c2ccc(Cl)cc2Cl)c1

TK198) Brc1ccc(NC2N(Cc3cccs3)C(=O)c3cccnc23)cc1

TK210) Cn1c2ccccc2c(O)c(I)c1=O

TK222) C1=CC=C2C(=C1)C(=O)NS2

NADase in vitro analysis of hSARM1 in the presence of 50µM NAD+ showed that these seven compounds have IC_50_ ≤ 10µM (Figure 3A and B, and figure 3 supplement 1). Control experiment with the known SARM1 inhibitor NAM (nicotinamide) showed IC_50_=43.3µM (Figure 3B), consistent with previously reported inhibitory values (Essuman et al., 2017). We next characterized the inhibitory mode of the following five compounds (TK 106, 138, 142, 210, 222) and measured the NADase activity at different NAD+ concentrations (1-160 µM) with and without the presence of 10µM of the five inhibitors. While TK106 showed a non-competitive inhibitory pattern, as seen in the Lineweaver-Burk plot, TK 138, 142, 210, and 222 showed a competitive pattern (Figure 3C).

To further characterize these inhibitors, we examined their species specificity and measured their effect on isolated zebrafish SARM1 (zfSARM1) compared to hSARM1. First, we measured the NADase activity and determined the 3D structure of purified zfSARM1 (Figure 4A-C). We found that the structure of zfSARM1 is very similar to that of hSARM1, with the latter’s atomic model docking very well into the former’s 3D cryo-EM reconstruction map (Figure 4A and B). Next, we measured the NADase kinetic behavior of zfSARM1 and found it to be similar to that of hSARM1, with Km and Kcat values of 33µM and 219min^−1^ for hSARM1 and 28µM and 411min^−1^ for zfSARM1 (Figure 4C). Like hSARM1, zfSARM1 also presents an apparent inhibition by substrate (as Ki for NAD+) and by product (as IC_50_ for NAM), with Ki=1350µM; IC_50_=227µM, compared to the hSARM1 Ki=727µM; IC_50_=43.3µM. Taken together, we have established that there is a high level of similarity in structure, regulation, and enzymatic activity between the human and zebrafish SARM1. However, we found that the responses of the zebrafish and human SARM1 towards the TK inhibitors, are in most cases inconsistent (Figure 4D). TK 142 and 222 - both Isothiazolinone derivatives, show a specific inhibitory effect on zfSARM1, albeit less clear at low inhibitor concentrations. On the contrary, TK138 has a specific activating effect, bringing up zfSARM1 NADase activity with EC_50_ (half-maximal effective concentration) at 2.7µM. Both the TK106 Calmidazolium and the TK210 Quinoline give an ambiguous response, that cannot be considered specifically inhibitory or activating.

**Figure 4:**
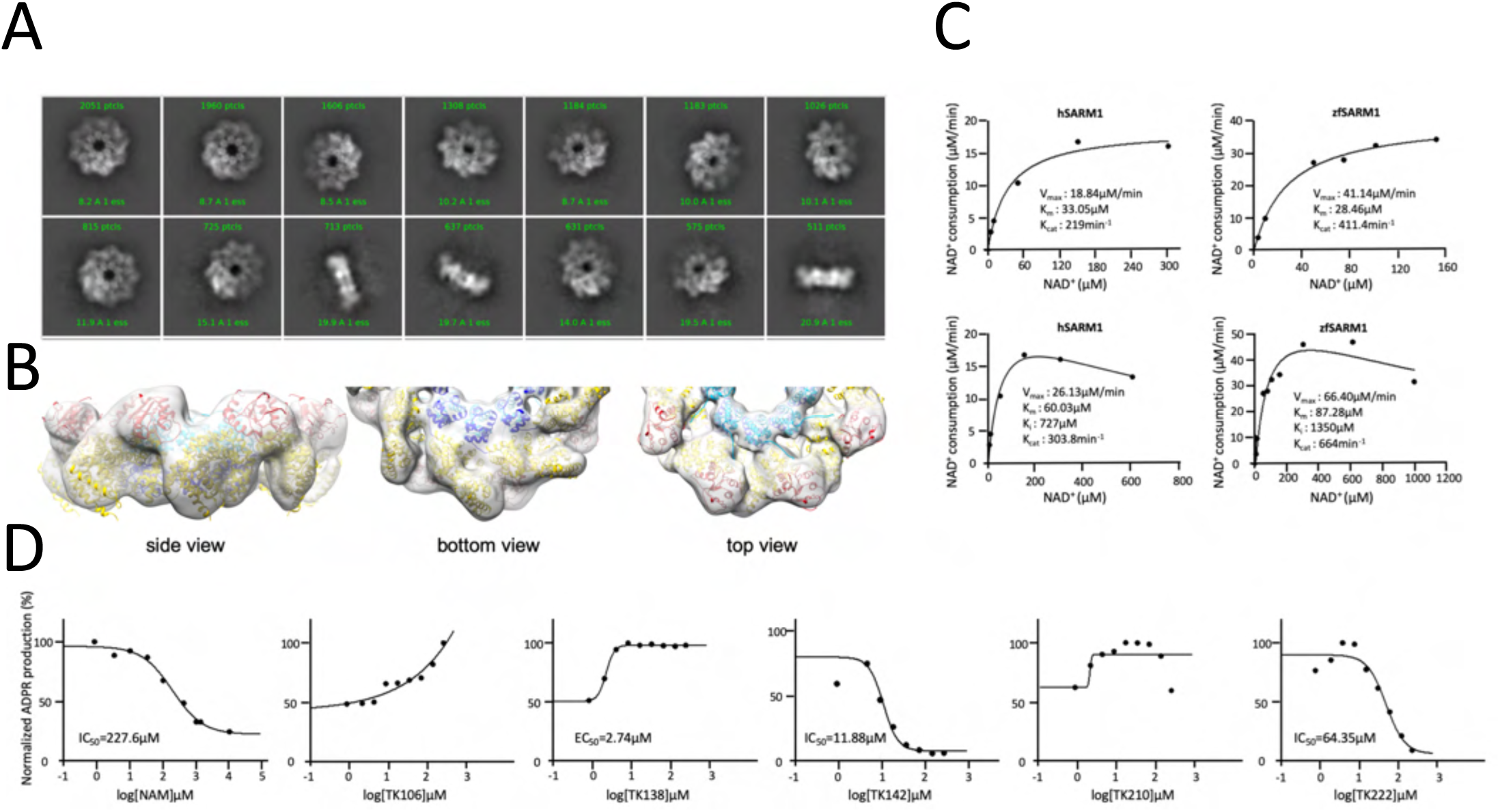
Structural, kinetic, and pharmacological characterization of the zebrafish (zf)SARM1. A) Selected representation of 2D class averages used for the 3D reconstruction of zfSARM1. The number of particles that were included in each class average are indicated. B) Color-coded (as in Figure 1A) protein model (of hSARM1 PDB ID 7ANW) docked in a transparent 8.1 Å resolution cryo-EM density map refined with C1 symmetry (gray). ‘Top view’ refers to the aspect of the molecule showing the TIR (red) and SAM2 (cyan) domains closest to the viewer, while in the ‘bottom view’ the SAM1 domains are the closest. C) Kinetic comparison between hSARM1 and zfSARM1 shows an overall similarity of the two, with close Km and Kcat values (upper panel). Inhibition by the substrate NAD+ is also observed in zfSARM1with a Ki of 1.3 mM, compared with a Ki of 0.7 mM for hSARM1 (lower panel). The kinetic parameters were determined from plots of reaction velocity of NAD+ consumption versus substrate (NAD+) concentration and then fitted to the Michaelis-Menten equation (Km and Vmax) or substrate inhibition equation (Ki) using non-linear curve fit in GraphPad Prism. Kcat was calculated by dividing the Vmax with protein molar concentration. D) The effect of hSARM1 inhibitors on zfSARM1 NADase activity. An IC_50_=227 µM for NAM demonstrates that zfSARM1 is also susceptible to inhibition by product, similar to hSARM1 (see figure 3B). While TK142 and TK222 exhibit concentration-dependent inhibitory effects on zfSARM1, it is not clear what are the effects of TK106 and TK210. Remarkably, TK138 has an opposite, activating effect, with an EC_50_ value of 2.7 µM. IC_50_ and EC_50_ values were determined according to ADPR production as measured in HPLC and calculated and plotted with GraphPad.

### Cryo-EM visualization of hSARM1 in the presence of the TK106 inhibitor

The kinetic Lineweaver-Burk analysis (Figure 3C) revealed a non-competitive mode of inhibition for TK106. Previously (Sporny et al., 2020), we discovered that large conformational changes in the peripheral ring (composed by the ARM and TIR domains) of the octamer correlate with hSARM1 NADase inhibitory (compact peripheral ring) and active (disordered peripheral ring) states. We also found that binding of NAD+ at the concave side of the ARM domain stabilizes the compact, inhibited conformation. To see whether TK106 inflicts a similar effect as NAD+ over the conformation of the peripheral ring, we visualized hSARM1 in the presence of 50µM of TK106 using cryo-EM.

For cryo-EM imaging, the near-intact hSARM1, lacking the N-terminal mitochondrial localization signal (^26^ERL...GPT^724^) and mutated in the NADase catalytic residue E642Q (hSARM1^E642Q^), was expressed in mammalian cell culture and isolated to homogeneity using consecutive metal chelate and size exclusion chromatography steps. We collected cryo-EM images of the purified hSARM1^E642Q^ - TK106 mixture. 2D classification revealed that about half of the particles in a face-view (where the ring hole is visible) were in the fully assembled 2-ring conformation (Figure 5 - figure supplement 1 A, B). This is despite the lack of NAD+, a condition that in our preparations normally shows a mere 10-20% of particles in that conformation (Li et al., 2021; Sporny et al., 2020). Remarkably, 2-D class averages of particles positioned in side-view revealed that 37% of them are arranged as duplexes of two hSARM1 ring octamers.

**Figure 5.**
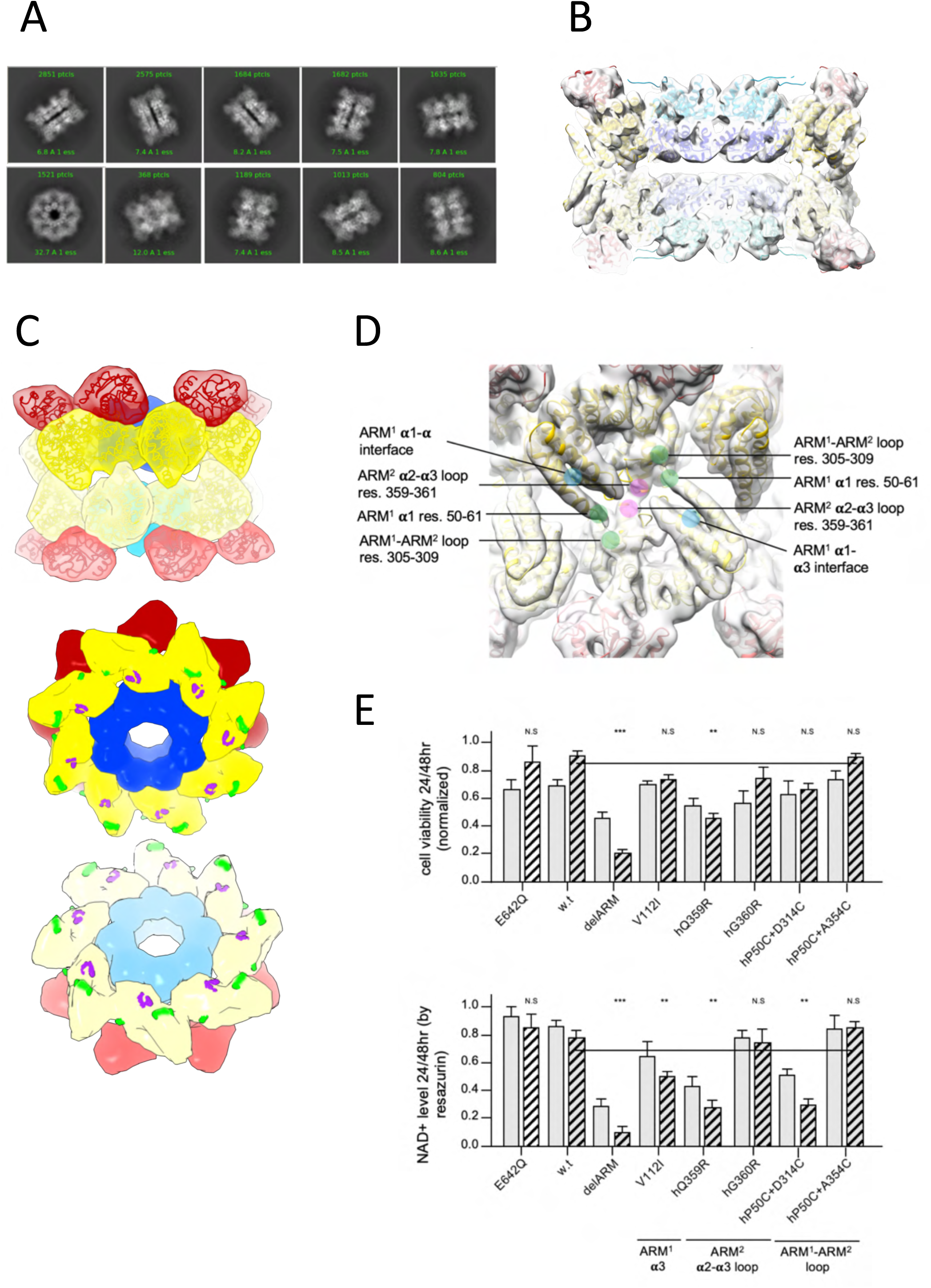
Cryo-EM structure of TK106 induced hSARM1 duplex. A) Selected representation of 2D class averages used for the 3D reconstruction. The number of particles included in each average is indicated at the top of each class. B) Protein model docked in a transparent 4 Å cryo-EM density map (gray), sliced at the frontal plane. Note that all the duplex interactions are mediated through the ARM domains (yellow), and none involves the SAM domains (blue). C) Top panel - surface representation of the hSARM1 duplex, color-coded as in (Figure 1A). The bottom monoplex is in lighter colors than the upper one. Bottom panel - ‘Open-book’ representation, where the two monoplexes are pooled apart to reveal the homotypic and heterotypic contacts points (colored as in Fig. 1 - supplement figure 1) on the ARM domains that mediate the duplex assembly. D) Zoom-in of the ARM-ARM contralateral duplex interactions. The interacting elements are colored and labeled. E) Cell toxicity (top panel) and NAD+ consumption (bottom panel) of the hSARM1 construct and mutants in HEK293F cells. The cells were transfected with hSARM1 expression vectors, as indicated. Cell viability counts and fluorescence resazurin NAD+ measurements were made 24 and 48h post-transfection. The NADase attenuated hSARM1^E642Q^ mutant was used as negative control. hSARM1^w.t.^ has very little NADase activity 48h post infection, while deletion of the inhibiting ARM domain (delARM) induces massive NAD+ consumption and cell death. Mutations (L112I, Q359R and P50C+D314C) at the duplex interaction sites of the ARM domain also induce NAD+ consumption (and less so cell death), however clearly milder than the fully active ‘delARM’ construct (three biological repeats, Student t test; *** p < 0.001; ** p < 0.05; n.s: no significance).

After several rounds of iterative 2D and 3D classification (Figure 5 - figure supplement 1B), we reconstructed 3D volume maps of the monoplex (single ring) and duplex forms of hSARM1 to 2.6 and 4.0 Å resolution, respectively (Figure 5 - and figure supplement 2). Overall, each of the two individual rings of the duplex, as well as the single monoplex ring, were very similar to our previously reported NAD+ supplemented structure (PDB code 7ANW) (Sporny et al., 2020). The map resolution (4.0 Å) falls short of revealing the binding site of TK106, but a strong density is visible at the concave surface of the ARM domains. Typically, in cryo-EM maps of hSARM1 not supplemented with NAD+, only very weak density occupies the concave ARM’s site (Figure 5 - figure supplement 3). This density could either account for a small molecule occupancy, like TK106, or rather be induced by the binding of TK106 at another site, which is difficult to trace.

The duplex is arranged as a back-to-back stack of two rings, where the contralateral SAM^1^ domains are close to each other, and the SAM^2^s are the furthest (Figure 5 B,C). However, the distance between the bottom parts of the SAM^1^ domains is about 10Å and unlikely to mediate any direct or indirect contacts. Instead, contralateral interactions are mediated solely by the ARM domains, via two sites (Figure 5 B-D). One interaction site is homotypic and involves the same ARM^2^ **α**2-**α**3 loop (res. QGK 359-361) from both ARM domains. The other interaction is heterotypic, where the N’ terminal region of ARM^1^ **α**1 helix from one protomer interacts with the upstream sequence PGRFA (res. 305-309) of the ARM^1^-ARM^2^ loop of the contralateral protomer, and vice versa (Figure 1 - figure supplement 1). This way, each ARM domain is not only held in place through lateral interactions with neighboring ARM, SAM, and TIR domains but also via contralateral interactions with the reciprocal SARM1 ring. These additional interactions are assumed to further stabilize the inhibitory conformation, preventing pre-mature activation.

### Mutagenesis of the hSARM1 duplex interface

To test this hypothesis, we introduced mutations designed to compromise the contralateral duplex ARM-ARM interactions. We have targeted the ARM^2^ **α**2-**α**3 loop (Q359R; G360R), the N’ terminal region of ARM^1^ **α**1 (P50C), the ARM^1^-ARM^2^ loop region (D314C, A354C), and the ARM^1^ 1-3 packing site (V112I), (Figure 5E). Q359 and G360 are located at the homotypic ARM^2^ **α**2-**α**3 loop. The two pairs of mutations - P50C+D314C, and P50C+A354C were introduced at the heterotypic site. V112I is likely to affect the packing and position of the ARM^1^ **α**1. It is a naturally occurring mutation recently found in ALS patients that was already tested and confirmed to induce mild elevated hSARM1 activity (Gilley et al., 2021). These constructs were transiently expressed in HEK-293F cells. The effect of these constructs on NAD+ levels and cell viability were monitored using a resazurin fluorescence assay and live cell counts. While G360R and P50C+A354C did not indicate for increased NADase activity, the V112I, Q359R and P50C+D314C mutants (Figure 5E) show a significant decrease in cellular NAD+ levels and (to a lesser extent) cell death in the course of the first two days after transfection. These NAD+ consumption and toxicity levels are milder than what was observed in our previous report (Sporny et al., 2020) with other ARM domain mutations. Such are the fully activated hSARM1 construct ‘delARM’ (res. 409–724) that is missing the entire ARM, as well as mutations that interfere with the inhibitory docking of TIR domains such as FP255-6RR, and mutations that obstruct NAD+ binding in the allosteric inhibitory site, such as W103D.

Taking together, the current and previous analysis of ARM mutagenesis leads to the conclusion that ARM auto-inhibition is mostly executed in the context of the monoplex ring, with duplex assembly that prevents NADase activity leakage.

### Axon degeneration assay - DRG axotomy

We next tested the selected inhibitors in the *in vitro* axonal degeneration assay (Shacham-Silverberg et al., 2018). In this assay, E13.5 mouse DRGs were plated and left to grow axons for 96 hours before severing the axons proximally to the DRG ganglion (Figure 6). While the severed axons from SARM1-/-mouse remained intact, as previously shown in (Osterloh et al., 2012), w.t DRG axons got fragmented and were eliminated almost completely 16 hours post axotomy. To test if our newly identified hSARM1 inhibitors can also protect axons from Wallerian degeneration, we supplemented the DRGs medium with 10-30µM of hSARM1 inhibitory compounds right before axotomy. To evaluate toxic side effects, we applied 10-30µM of the inhibitory compounds to DRG explants without performing axotomy. Overall, toxicity was manifested as the thinning, and in some cases elimination, of axons’ extremities, at the growth cones region. However, other toxic effects were also observed, such as the elimination of the segments of axons proximal to the DRG ganglion (e.g. by TK106). Mild manifestations of such toxic effects were observed for almost all tested compounds, usually in a concentration-depended manner. However, only 20 µM of TK198 were toxic in a statistically significant manner. The most effective compounds in providing a high level of protection while being only very mildly toxic are TK210, TK138, TK222, and to a lesser extent, TK106. TK142 is not toxic at all but provides only mild protection. In this assay, TK198 and TK174 did not protect axons after axotomy, and while TK198 was toxic, TK174 was not. It is possible that TK174 and TK142 are neither toxic nor protective due to possible impermeability of these compounds into the neurons’ cytosol.

**Figure 6:**
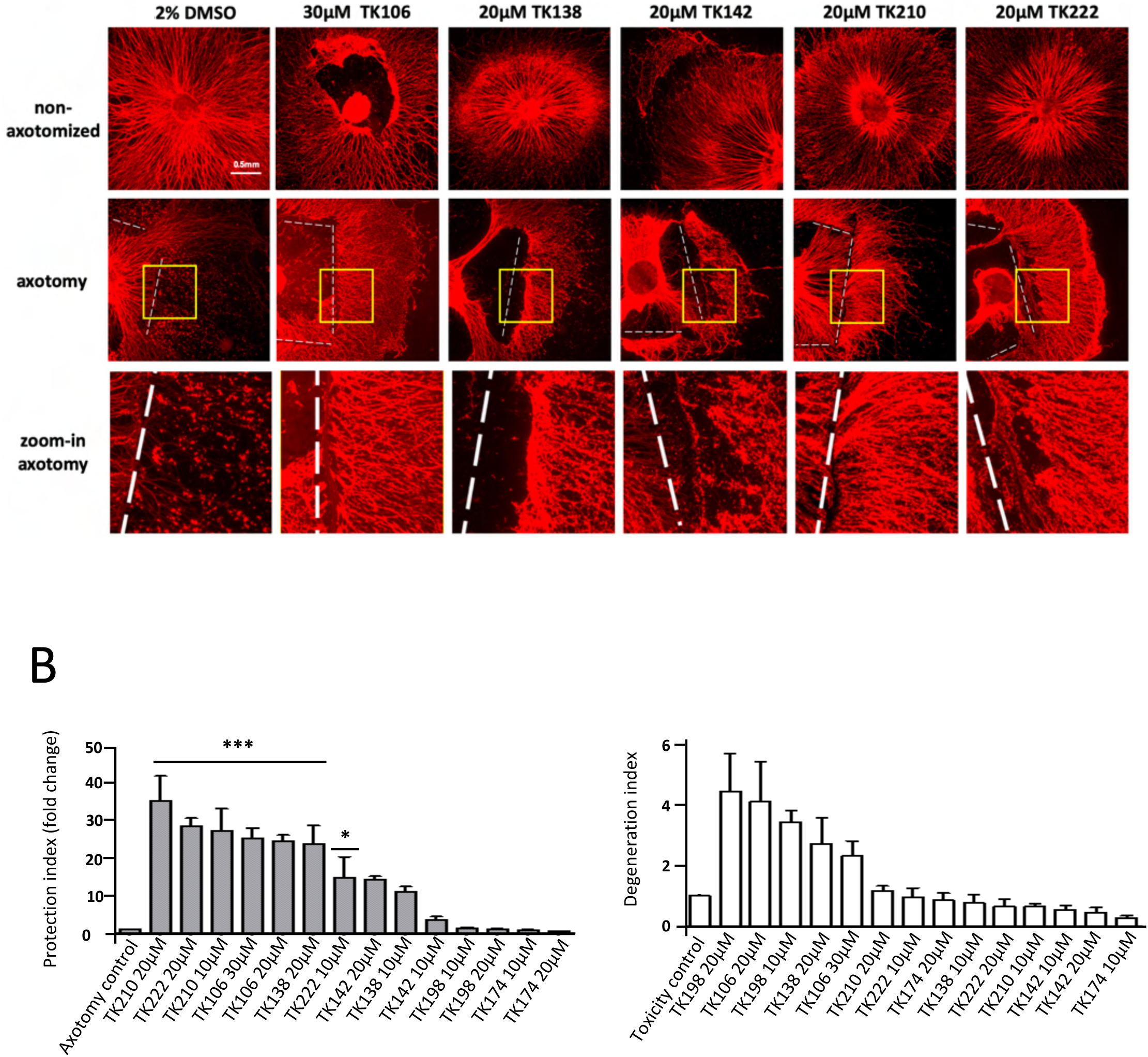
hSARM1 inhibitory compounds protects axons against axotomy-induced degeneration. A) Mouse DRG explants cultures were grown in NGF-containing media for 96 hours. Right before axotomy, the media was exchanged and supplemented with either 10-30µM of the hSARM1 inhibitory compounds or just with 0.2% DMSO for control. Then the cultures were left to grow for additional 16 hours. The white dashed lines indicate cut site (axotomy). To evaluate toxic side effects, we applied 20-30µM of the inhibitory compounds without performing axotomy. The axons were stained with anti-Tuj1 antibody. B) Quantification of the protective (left) and toxic (right) effects induced by the hSARM1 inhibitory compounds. While TK210, TK106, and TK138 provide the highest level of protection after axotomy. TK106 (and to a lesser extent TK138) has a significant toxic effect also without axotomy. Data are presented as mean ± SEM for 3 independent experiments. One-way ANOVA followed by Dunett post-tests were used to determine statistical significance (*p<0.05, **p<0.01, ***p<0.0001). All the analyses were performed in GraphPad Prism software.

## Discussion

Some aspects of SARM1 regulation, inhibition, and activation are still unclear. For example, there are gaps in our understanding of the allosteric regulation of SARM1, where NAD+ is thought to prevent activation - and NMN to induce it, reflecting the changes in the cellular levels of these metabolites in homeostasis and under stress. However, the concentrations in which these and other nicotine nucleotide compounds affect SARM1 activity in vitro are significantly higher than their in vivo levels. We have previously observed (Sporny et al., 2020) that a supplement of 5mM NAD+ enforces the autoinhibited, ordered, conformation of the peripheral ARM-TIR ring, which is otherwise disordered in most hSARM1 particles in our cryo-EM samples. Also, 2mM NAD+ significantly decreases the rate of NADase activity when compared to commercial porcine brain NADase. However, the physiological levels of NAD+ are thought to be much lower, ranging from 0.1 to 0.8mM (Cambronne et al., 2016; Hara et al., 2019; Houtkooper et al., 2010; Liu et al., 2018b). Considering that the calculated substrate inhibition constant Ki is 0.7mM (this work, Figure 4C) or 0.3mM (Angeletti et al., 2021), NAD+ alone would be insufficient in maintaining effective autoinhibition. We therefore propose that while NAD+ is not able to hold tight inhibition alone, accumulative inhibition together with NAM (competitive) and NaMN (allosteric), as well as other molecules currently unknown to us, keep the catalytic activity of SARM1 below a degenerative-inducing level. Also, we present here the possibility for a more inhibited form of SARM1 as presented in the cryo-EM structure of the TK106 - supplemented hSARM1 (Figure 5). It is a meta-stable duplex structure in which each protomer is held in place not only by lateral interactions but also via contralateral contacts that would make disassembly and NADase activity much harder. Indeed, site-directed mutagenesis at the duplex interaction sites results in significant elevation of NADase activity in cultured cells (Figure 5E). Maybe the most interesting of these activating mutations is the substitution of V112 to isoleucine, which was also recently identified as a SARM1 activating mutation in ALS patients (Gilley et al., 2021). V112 is located at ARM^1^ **α**3 facing **α**1, and it is difficult to understand how can such a mild mutation at the most distal part of the ARM domain, which is not engaged in any domain-domain interactions in the monoplex structure, inflict elevation of NADase activity. Only when considering the role of **α**1 in the duplex assembly (Figure 5D), one realizes that even a slight rearrangement can lead to duplex dissociation and NADase activation.

Another unclear aspect in SARM1 research is the NADase catalytic activity of the TIR domain. For instance, there is no detailed structural information of active “caught in the action” SARM1 exists, and data are limited to cryo-EM and crystal structures of TIR without NAD+. It is possible that we are still missing important information about the mechanism of TIR NAD+ catalysis and that other, yet unknown, sites that regulate catalysis exist. Such alternative regulatory mechanisms are implied from the crystal packing of the x-ray crystal structure of hSARM1 TIR domain (Horsefield et al., 2019), as well as the related plant TIR-based ‘resistome’ structures (Ma et al., 2020; Martin et al., 2020) and the bacterial BcPycTIR anti phage system (Tal et al., 2021). Our data also indicate additional regulatory sites. We found that TK138 is a potent (IC_50_=2.9µM) competitive inhibitor of hSARM1 (Figure 3), implying a direct engagement with the NADase active site. However, in zfSARM1, TK138 activates the NADase activity with EC_50_=2.7µM (Figure 4D). Considering the similarity between zfSARM1 and hSARM1 in structure (Figure 4 A and B), kinetic values (Figure 4C), and their common inhibition by substrate (NAD+) and product (NAM), it is likely that TK138 affects the same active site as in hSARM1, only with opposite outcome in zfSARM1. This means that small molecules, such as TK138, can increase the NADase catalytic activity by binding to the active site and not only to a distal allosteric site like the ARM domain.

In summary, in this work, we present new data that expand the molecular perspective of SARM1 activation and auto-inhibition. So far, we presumed that the tight assembly of the SARM1 octamer preserves full autoinhibition and that the disassembly of the peripheral ARM-TIR ring allows maximal activation. However, analyzing the kinetic and structural effects of several hSARM1 inhibitors that we here discovered, shows that activation can be further augmented following the binding of small molecules to the TIR domain, and that autoinhibition can be strengthen by the formation of a meta-stable duplex structure, which has not been described before. Altogether, the discovered inhibitory compounds and the new mechanistic realizations that these inhibitors revealed about SARM1 regulation will help develop safer and more effective drugs to treat SARM1-induced neuropathies.

## Materials and methods

### Key Resources Table

| Key Resources Table |  |  |  |  |
| --- | --- | --- | --- | --- |
| Reagent type, species, or resource | Designation | Source or reference | Identifiers | Additional information |
| gene (homo sapiens) | SARM1 | Imagene | uniprot Q6SZW1 |  |
| cell line (homo sapiens) | HEK293F | Thermo Fisher Scientific | Cat# R79007 |  |
| cell line (homo sapiens) | HEK293T | ATCC | CRL-11268 |  |
| gene (Zebrafish) | SARM1 | Imagene | uniprot F1QWA8 |  |
| transfected construct (mammalian vector) | pEGFP-N1 | Clontech |  | modified |
| chemical compound, drug | Polyethyleneimine 40 kDa | Polysciences | PEI-MAX |  |
| chemical compound, drug | Resazurin sodium salt | SIGMA | R7017 |  |
| chemical compound, drug | NAD<sup>+</sup> | SIGMA | NAD 100-RO |  |
| chemical compound, drug | FMN | SIGMA | F8399 |  |
| chemical compound, drug | ADH | SIGMA | A3263 |  |
| chemical compound, drug | Diaphorase | SIGMA | D5540 |  |
| chemical compound, drug | BSA | ORNAT | B9001S |  |
| chemical compound, drug | eNAD | SIGMA | N2630 |  |
| chemical compound, drug | DMSO | SIGMA | 67-68-5 |  |
| chemical compound, drug | TK222 | SIGMA | 561487 |  |
| chemical compound, drug | TK106 | MolPort | MolPort-002-913-512 |  |
| chemical compound, drug | TK138 | MolPort | MolPort-023-276-485 |  |
| chemical compound, drug | TK142 | MolPort | MolPort-002-903-305 |  |
| chemical compound, drug | TK210 | MolPort | MolPort-002-044-160 |  |
| chemical compound, drug | TK174 | MolPort | MolPort-019-885-142 |  |
| chemical compound, drug | TK198 | MolPort | MolPort-007-649-343 |  |
| mice | DRG | Envigo Israel | CD(ICR) |  |
| Cell culture slide | 8-chamber | SPL Life Sciences | 30108 |  |
| chemical compound, drug | poly-d-lysin | Sigma aldrich | P6407 |  |
| chemical compound, drug | Laminin | Sigma aldrich | L2020 |  |
| chemical compound, drug | Neurobasal™-A | GIBCO | 10888022 |  |
| chemical compound, drug | B-27 | GIBCO | 17504044 |  |
| chemical compound, drug | penicillin-streptomycin | Biological Industries | 03-031-1B |  |
| chemical compound, drug | L-glutamine | Biological Industries | 03-020-1B |  |
| chemical compound, drug | NGF | Alomone labs | mNGF 2.5 S N-100 |  |
| chemical compound, drug | Formaldehyde 4% | Bio-Lab | 6450323F1 |  |
| Antibody | anti-tubulin b III antibody (Tuj1 clone) | R&D systems | MAB1195 |  |
| Antibody | Cy3 goat anti-mouse | Jackson ImmunoResearch Laboratories | 115-165-146 |  |
| microscope | Eclipse Ni-E upright microscope system | Nikon |  |  |
| software, algorithm | MotionCorr2 | (Li et al., 2013) |  |  |
| software, algorithm | Gctf | (Zhang, 2016) |  |  |
| software, algorithm | Cryosparc V2 | (Punjani et al., 2017) |  |  |
| software, algorithm | SCIPION wrapper | (Martinez et al., 2020) |  |  |
| software, algorithm | Warp | (Tegunov and Cramer, 2019) |  |  |
| software, algorithm | CCP4 | (Winn et al., 2011) |  |  |
| software, algorithm | GESAMT | (Krissinel, 2012) |  |  |
| software, algorithm | MOLREP | (Vagin and Teplyaev, 2010) |  |  |
| software, algorithm | Coot | (Emsley et al., 2010) |  |  |
| software, algorithm | REFMAC5 | (Murshudov et al., 2011) |  |  |
| software, algorithm | JLIGAND | (Lebedev et al., 2012) |  |  |

### cDNA generation and subcloning

Cloning of all the constructs was made as detailed in (Sporny et al., 2020) by PCR amplification from the complete cDNA clone (Imagene) of hSARM1 (uniprot: Q6SZW1) and of Zebrafish (uniport: F1QWA8). For expression in mammalian cell culture, the near-intact hSARM1^w.t.^ (^26^ERL...GPT^724^), the mutant hSARM1^E642Q^ and zfSARM1^w.t^ (^22^DRL...QKK^713^) constructs were ligated into a modified pEGFP-N1 mammalian expression plasmid which is missing the C-terminus GFP fusion protein, and includes N-terminal 6*HIS-Tag followed by a TEV digestion sequence.

### Protein expression and purification

Expression and purification were made as detailed in (Sporny et al., 2020). For protein purification, hSARM1^w.t.^ hSARM1^E642Q^ and zfSARM1^w.t.^ were expressed in HEK293F suspension cell culture, grown in FreeStyle™ 293 medium (GIBCO), at 37°C and in 8% CO_2_. Transfection was carried out using preheated (70°C) 40 kDa polyethyleneimine (PEI-MAX) (Polysciences) at 1mg of plasmid DNA per 1 liter of culture once cell density has reached 1*10^6^ cells/ml. Cells were harvested 4- (in the case of hSARM1^w.t.^ and zfSARM1^w.t.^) and 5- (in the case of hSARM1 ^E642Q^) days after transfection by centrifugation (10 min, 1500 x g, 4°C), re-suspended with buffer A (50mM Phosphate buffer pH 8, 400mM NaCl, 5% glycerol, 1mM DTT, 0.5mM EDTA, protease inhibitor cocktail from Roche) and lysed using a microfluidizer followed by two cycles of centrifugation (12000 x g 20 min). Supernatant was then filtered with a 45µm filter and loaded onto a pre-equilibrated Ni-chelate column as in (Guez-Haddad et al., 2015; Yom-Tov et al., 2017). The column was washed with buffer A supplemented with 25mM Imidazole until a stable baseline was achieved. Elution was then carried out in one step of 175mM Imidazole, after which protein-containing fractions were pooled and loaded onto pre-equilibrated Superdex 200 Increase 10/300 (GE Healthcare) for size exclusion chromatography. The elution was performed with 25mM Phosphate buffer pH 8.5, 120 mM NaCl, 2.5% glycerol, and 1mM DTT. Protein-containing fractions were pooled and concentrated using a spin concentrator to 1.5 mg/ml. The concentrated proteins were split into aliquots, flash-frozen in liquid N_2_ and stored at −80°C for later cryo-EM visualization and enzymatic assays.

### High-throughput inhibitor screening of potential hSARM1 inhibitors

The high-throughput inhibitor-screening assay was performed using the MultiDrop 384 microplate dispenser (Thermo Scientific) and 1536-well black/black non-binding Microplates (Greiner Bio-One). A total of ~150,000 compounds from the following libraries were screened: LOPAC/SIGMA (Navigator LOPAC^1280^) 1280 compounds; MicroSource (Spectrum Collection-Known Drugs (66%), Natural Products (26%) and Other Bioactive Components (8%)) 2400 compounds; Prestwick (Prestwick Chemical Library® - approved drugs) 1200 compounds; New Selleck collection 2020 (Bioactive) 3727 compounds; GSK Inhibitors (GSK Published Kinase Inhibitor Set (PKIS)) 367 compounds; Analyticon (MEGxp: Pure natural compounds from plants) 436 compounds; Analyticon (MEGxm: Pure natural compounds from microorganisms) 469 compounds; Analyticon (MEGxp: Pure natural compounds from plants) 3540 compounds; Selleck Chemicals (Natural Product Library) 144; Enamine (Drug-Like Set (DLS)) 20160 compounds; MayBridge (HitFinderTM Collection) 14400 compounds; ChemDiv (DIVERSet) 100000 compounds; Kinom Set (Kinom Set) 187 compounds; SGC (Epigenetic chemical probes) 97 compounds; FDA cancer library (Anti-cancer FDA approved drugs) 248 compounds.

First, 10 nL of each screened compound, solubilized in DMSO to 10mM, were plated using Echo® 550 Liquid Handler (Sunnyvale, CA, USA). Then, 5 µl of purified hSARM1^w.t.^ (at 50 nM concentration) in 25 mM HEPES pH 7.5, 150mM NaCl were added to each well for 10 minutes incubation at 25°C. Next, 2.5 µl of NAD+ (0.25 µM, in 25 mM HEPES pH 7.5, 150mM NaCl) were added to each well and incubated for 70 min at 37°C. The compounds concentration at this point was 13.3 µM.

Measurement of NAD+ concentrations was made by a modified enzymatic coupled cycling assay. 2.5 µl of the reaction mix, which includes 100 mM Phosphate buffer pH=8, 0.195% ethanol, 1 µM FMN (Riboflavin 5’-monophosphate sodium salt hydrate), 6.8 U/ml Alcohol dehydrogenase (SIGMA), 0.45 U/ml Diaphorase (SIGMA), 1.25 mg/ml BSA (SIGMA), 1.25% Tween 20 (SIGMA) and freshly dissolved (in DDW) 2 µM Resasurin (SIGMA), was added to each well plate and incubated in dark for 1h to 3h at 25°C. Fluorescent signals were measured by PheraStar FS plate reader (BMG Labtech, Ortenberg, Germany) at 540-nm excitation and 590-nm emission wavelengths. Data analysis was performed using Genedata software. All values were normalized to positive (no hSARM126-724) and negative (with hSARM126-724) internal controls per plate.

### Small molecules QC Result analysis

The identity PASS criteria are when a protonated molecular ion, adduct, or simple fragment are found with relative abundancy > 70 % e.g., M-H, M+H, M+Na, M+DMSO, M+H +ACN, M-H + formic acid. The purity PASS criteria are when (i) the signal from the Photo Diode Array PDA detector is greater than or equal to 70% of the expected signal, or (ii) ESLD 2 70% with no PDA, or (iii) PDA 2 70% and ELSD 2 70%. The identity FAIL criteria are when the expected mass ions are not found by positive or negative ionization mode using ESI or APCI methods. The purity FAIL criteria are when (i) PDA < 70% with no ELSD, or (ii) ELSD < 70% with no PDA, or (iii) PDA < 70% and ELSD < 70%. A sample must meet both identity and purity PASS criteria in order to get a QC Result of PASS. If results are ambiguous; for example, due to a method failure or the absence of the required ionization method, then the sample gets a QC Result of TENTATIVE.

Samples were dissolved in (50 µL, 60 µM) 384-well plates in water: acetonitrile (7:3) trace DMSO; were injected (7µL) onto an Acquity UPLC® BEH C18 1.7 µm 2.1×50 mm Column at a flow rate of 0.5 mL/min. The mobile phase solvents were as follows: A = 0.05% Formic Acid in water : acetonitrile (95:5), B = 0.05% Formic Acid in acetonitrile. Elution was achieved via the delivery of a 1 min hold at 100% A 3 min 0-100% B gradient followed by a 1.0 min hold at 100% B prior to re-equilibration. Instrument Specifications: Waters Acquity UPLC® H class with PDA detector, ELSD and using Acquity UPLC® BEH C18 1.7 µm 2.1×50 mm Column (PN:186002350, SN 02703533825836). MS-system: Waters, SQ detector 2

### NADase HPLC analysis

For validating the first round of potential inhibitors, purified hSARM1^w.t.^ was first diluted to 125nM in 25mM HEPES pH 7.5, 150mM NaCl, and incubated for 10 minutes in 25°C with 10 µM of each inhibitor compound. Then, 50µM of NAD+ (in the same buffer) in a 1:1 v/v ratio were added and incubated for 30 min in 37°C. 1:100 (v/v) BSA (NEB Inc. 20mg/ml) was included, and reactions were stopped by heating at 95°C for 2 minutes and centrifuged.

HPLC measurements were performed using a Merck Hitachi Elite LaChrom HPLC system equipped with an autosampler, UV detector and quaternary pump. HPLC traces were monitored at 260nm and integrated using EZChrom Elite software. 10 µL of each sample were injected onto a Waters Spherisorb ODS1 C18 RP HPLC Column (5 µm particle size, 4.6 mm x 150 mm ID). HPLC solvents are; A: 100% methanol; B: 120mM sodium phosphate pH 6.0; C double-distilled water (DDW). The column was pre-equilibrated with B:C mixture ratio of 80:20. Chromatography was performed at room temperature with a flow rate of 1.5 ml/min. Each analysis cycle was 12 min long as follows (A:B:C, v/v): fixed 0:80:20 0-4 min; gradient to 20:80:0 4-6 min; fixed 20:80:0 from 6-9 min, gradient to 0:80:20 from 9-10 min; fixed 0:80:20 from 10-12 min. The NAD+ hydrolysis product ADPR was eluted at the isocratic stage of the chromatography while NAD+ elutes in the methanol gradient stage.

Compounds that inhibit at least 50% of NADase activity at 10µM, were further examined for half maximal inhibitory concentration (IC_50_), where hSARM1^w.t.^ and zfSARM1^w.t.^ were pre-incubated with doses of inhibitor compounds for 10 min at room temperature −25°C, before the NADase reaction was commenced.

### Calculation of SARM1 kinetic parameters

For *V*_max_ and *K*_m_ determination, the NADase activity assay was performed with several different NAD+ substrate concentrations and sampled in constant time points. For each NAD+ concentration, a linear increase zone was taken for slope (V_0_) calculation. All data were then fitted to the Michaelis-Menten equation using non-linear curve fit by graphpad. Vmax was determined and *K*_cat_ was calculated by dividing the *V*_max_ with protein molar concentration. *K*_cat_ was calculated by dividing the *V*_max_ with protein molar concentration. Lineweaver-Burk plots of (1/V) versus 1/[S] in the presence of a constant concentration of inhibitors were used to determine the type of the enzyme inhibition. *Determination of IC*_*50*_

#### Determination of IC_50_

Serial concentrations of inhibitor compounds were pre-incubated with 0.33 µM hSARM1 or 0.35 µM zfSARM1 for 10 minutes at room temperature, followed by the addition of 50 µM NAD. After incubation of 10 minutes at 37°C, the reactions were stopped by heating at 95°C for 2 minutes. The amount of ADPR product was measured by HPLC and used to calculate the initial rate. IC_50_ values were calculated by plotting the initial rate to the dose of the compounds (log10).

All calculations were performed in GraphPad Prism software.

### NADase activity in cultured cells assay

HEK293F cells were seeded in 24-well plates (1 million cells in each well) in a final volume of 1 ml of FreeStyleTM 293 medium (GIBCO). The cells were transfected with 1 µg DNA of different hSARM1 constructs and incubated at 37°C and in 8% CO2. After 24hr, 100µl of cells were collected from each well every day and centrifuged at 600g for 10 min at room temperature. The pelleted cells were then resuspended with 0.03mg/ml Resazurin sodium salt (SIGMA) dissolved in FreeStyleTM 293 medium (GIBCO). All the samples were incubated for 45 min at 37°C, centrifuged (10min, 600g, 25°C) and then transferred to a 384-well black plate (Corning). Fluorescent data were measured using a SynergyHI (BioTek) plate reader at 560 nm excitation and 590 nm emission wavelengths. All fluorescent emission readings were averaged and normalized by subtracting the Resazurin background (measured in wells without cells). Also, live cell numbers and present from total cells were counted using the trypan blue viability assay and cell counter.

### Axon degeneration assay after axotomy

#### Explant culture

Dorsal root ganglion (DRG) were dissected from E13.5 wild-type mice and plated in a 8-wells chamber coated with 10µg/ml poly-d-lysin (Sigma-Aldrich, St Louis, MO) and 10µg/ml mouse Laminin (Sigma-Aldrich). The DRGs were grown in Neurobasal™-A (NB, GIBCO, 10888022) medium supplemented with B27 (Gibco, Waltham, MA), penicillin-streptomycin solution, glutamine (Biological Industries, Beit HaEmek, Israel), and NGF (Alomone Labs, Jerusalem, Israel) in a humidified incubator (37°C, 5% CO2).

#### SARM1 inhibitors treatment and axotomy

After 96h, the medium was exchanged to NGF containing medium, supplemented with 10-30µM SARM1 inhibitory compounds, which were first dissolved in DMSO to a 10mM stock concentration. For axotomy, axons were cut using a needle, in close proximity to the cell body and were examined 16 hours post-axotomy.

#### Immunohistochemistry

DRG cultures were fixed in 4% formaldehyde (Bio-Lab) and 15% sucrose in PBS for 1hr and gently washed 3 times with PBS. Cultures were blocked in blocking solution (3% BSA and 0.1% Triton X-100 in PBS) for 1 hour and then incubated overnight in primary antibody Tuj-1 at 4°C. Samples were gently washed 3 times with PBS and incubated with second antibody goat anti-mouse Cy3 for 1 hour at room temperature and finally washed 3X in PBS.

#### Quantification of axonal protection and degeneration

In situ images of DRG explants were taken with Eclipse Ni-E upright Nikon microscope with DS-Qi2 Monochrome cooled digital camera.

Images were binarized such that pixels corresponding to axons converted to white, while all other regions converted to black. The images were analyzed using ImageJ software with algorithm that distinguishes fragmented from intact axonal segments. To each image we calculated the degeneration index (DI) defined as the ratio of fragmented to intact axon number. To quantify axonal protection, we normalized the DI values to DMSO with axotomy, and then we divide one by each calculated value. The data represented as fold change between axotomy with treatments of SARM1 inhibitory compounds to axotomy without treatments. To quantify axonal degeneration caused by the compounds’ toxicity, we normalized the DI values to DMSO control without axotomy and generated a degeneration index. From each explant, 4 nonoverlapping frames were randomly collected and quantified per experimental condition.

### Cryo-EM grids preparation

Cryo-EM grids were prepared by applying 3 µl protein samples to glow-discharged (PELCO easiGlow™ Ted Pella Inc., at 15 mA for 1 minute) holey carbon grids (Quantifoil R 1.2/1.3, Micro Tools GmbH, Germany). The grids were blotted for 4 seconds and vitrified by rapidly plunging into liquid ethane at −182 °C using Leica EM GP plunger (Leica Microsystems, Vienna, Austria). The frozen grids were stored in liquid nitrogen until the day of cryo-EM data collection.

### Cryo-EM data acquisition and processing

Talos Glacios in EMBL, Grenoble, France was used for grid screening in preparation of Titan Krios data collection in ESRF (CM01). Frozen grids were loaded onto a 200kV Talos Glacios (ThermoFisher) electron microscope equipped with a Falcon3 direct electron-counting camera (ThermoFisher). Cryo-EM data were acquired with EPU software (FEI) at a nominal magnification of 120,000, with a pixel size of 1.224A, in linear mode and a total dose of ~44 electrons per Å2. A dataset of zebrafish SARM1 (w/o NAD) (6006 micrographs) was collected. Preprocessing was performed in Warp pipeline including motion correction, CTF estimation and particle picking (Langer et al., 2008). Further data processing was conducted using the cryoSPARC suite, including 2D classification, ab-initio reconstruction and refinement. A total of 885116 particles were imported, from which 52259 particles were selected based on iterative reference-free 2D classifications for reconstruction. Initial maps were calculated using ab-initio reconstruction and higher-resolution maps were obtained by imposing C8-symmetry in non-uniform 3D refinement. Working maps were locally filtered based on local resolution estimates.

Titan Krios in ESRF CM01 beamline (Kandiah et al., 2019) at Grenoble, France, was used for data collection of the TK106 supplemented hSARM1^E642Q^ (10761 micrographs) and zebrafish SARM1 (+NAD) (6298 micrographs). Frozen grids were loaded into a 300kV Titan Krios (ThermoFisher) electron microscope (CM01 beamline at ESRF) equipped with a Gatan K3 direct electron detector and a Gatan Bioquantum LS/967 energy filter. Cryo-EM data were acquired with EPU software (FEI) at a nominal magnification of x105,000, with a pixel size of 0.839 Å. 10761 movies of the TK106 supplemented grid sample were acquired in superresolution mode at a flux of 14.855 electrons per p^2^ s^−1^, giving a total exposure of ~41 (zf SARM1) or ~42 electrons (TK106 supplemented hSARM1^E642Q^) per Å^2^ and fractioned into 40 frames. A defocus range from −0.8µm to −2.8 µm were used. Using the SCIPION wrapper (Martinez et al., 2020) the imported movies were drift-corrected using MotionCor2 and CTF parameters were estimated using Gctf for real-time evaluation. In SCIPION wrapper, an initial set of particles were picked by crYOLO autopicking, extracted and classified using Relion. The motion-corrected micrographs were re-evaluated for CTF parameters re-picked in Warp. Further data processing was conducted using the cryoSPARC suite. In case of TK106 supplemented hSARM1^E642Q^, a total of 1420569 particles were imported. Details of image processing can be found in Figure 5 supplementary figure 1. 2D classification yielded, 519892 particles containing inner + outer rings or duplexes. 478,523 particles contained only inner rings. Initial maps were calculated using ab-initio reconstruction and higher-resolution maps were obtained by imposing C8-symmetry in non-uniform 3D refinement. Working maps were locally filtered based on local resolution estimates. Reference based repicking in CryoSPARC suite yielded 2193205 particles and a better result in terms of the monoplex maps. Two final maps were derived, one containing a monoplex at 2.57 A (~998000 particles) and the second map contained a duplex at 4.02 A (~28000 particles).

In case of zebrafish SARM1 (+NAD), a total of 671383 particles were imported. 211482 particles contained inner + outer rings. Initial maps were calculated using ab-initio reconstruction and higher-resolution maps were obtained by imposing C8-symmetry in non-uniform 3D refinement. Working maps were locally filtered based on local resolution estimates.

### Model building and refinement

We used our previously determined hSARM1 octamer structure (PDB 7ANW) as template in docking into the duplex map, and truncated regions where no clear density is observed. Refinement was performed using PHENIX (Liebschner et al., 2019) RealSpaceRefine (Afonine et al., 2018) program.

### Accession numbers

Coordinates and map have been deposited in the Protein Data Bank and in the EMDB with the following accession codes: PDB ID 7QG0, EMD-13951

## Acknowledgments

We acknowledge the European Synchrotron Radiation Facility for provision of beam time on CM01 and thank the staff of beamline CM01 of ESRF and members of the Opatowsky lab for technical assistance. We thank Gershon Kunin for IT management. The Israel National Center for Personalized Medicine is supported by a research grant from the Nancy and Stephen Grand. This work was supported by funds from ISF grants no. 1425/15 and 909/19 and BSF grant no. 2019150 to Y.O. Y.O. is a Katzir Professorial Chair of Biophysics, and A.Y. is an incumbent of the Jack and Simon Djanogly Professorial Chair in Biochemistry.

## Figure legends

**Figure 1 supplement 1.**
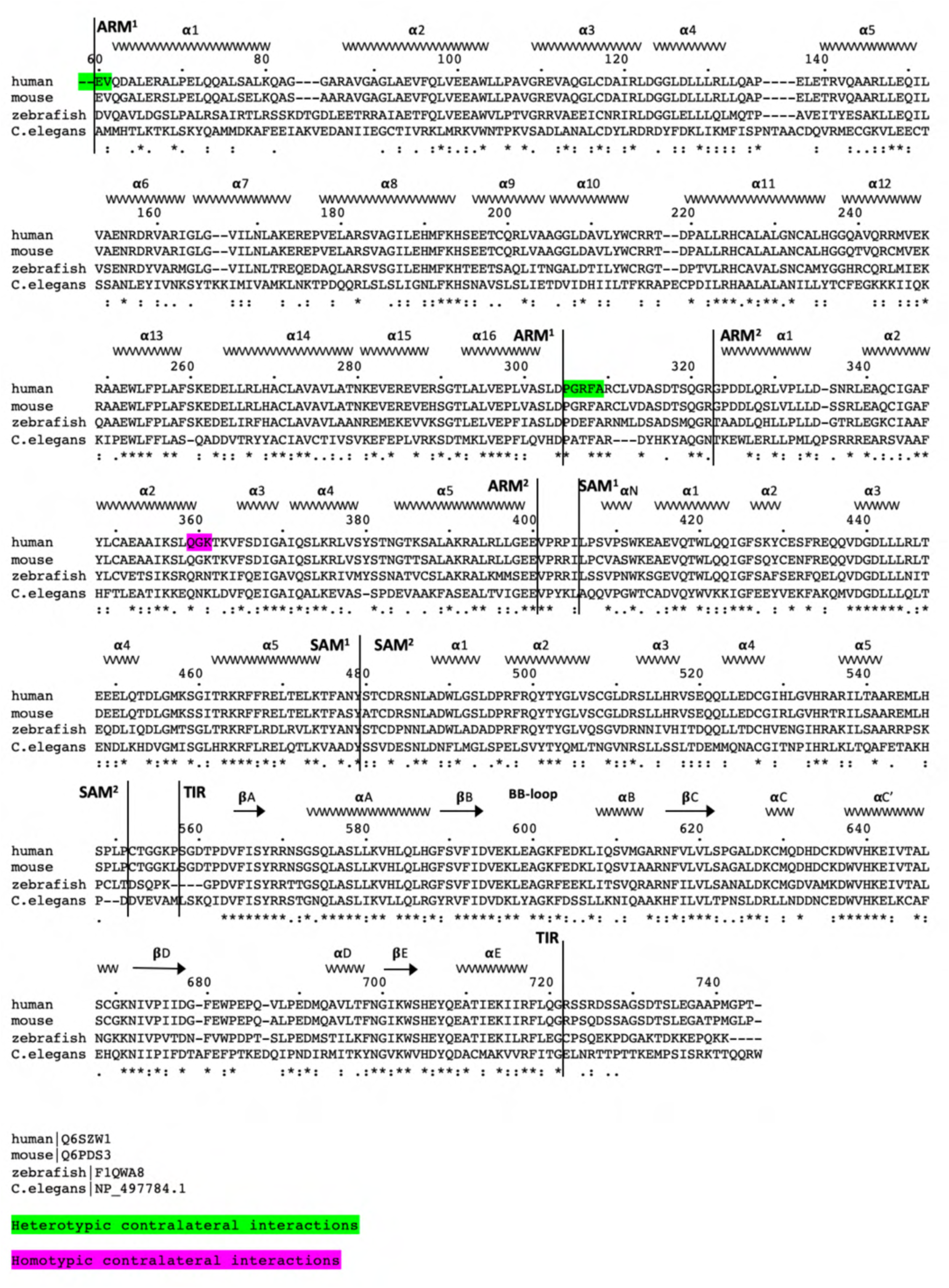
Structure-based sequence alignment of the SARM1 of human, mouse, zebrafish, and the C. elegans homolog TIR-1. Color-coded highlights and Uniprot protein accession numbers are listed below.

**Figure 3 supplement 1.**
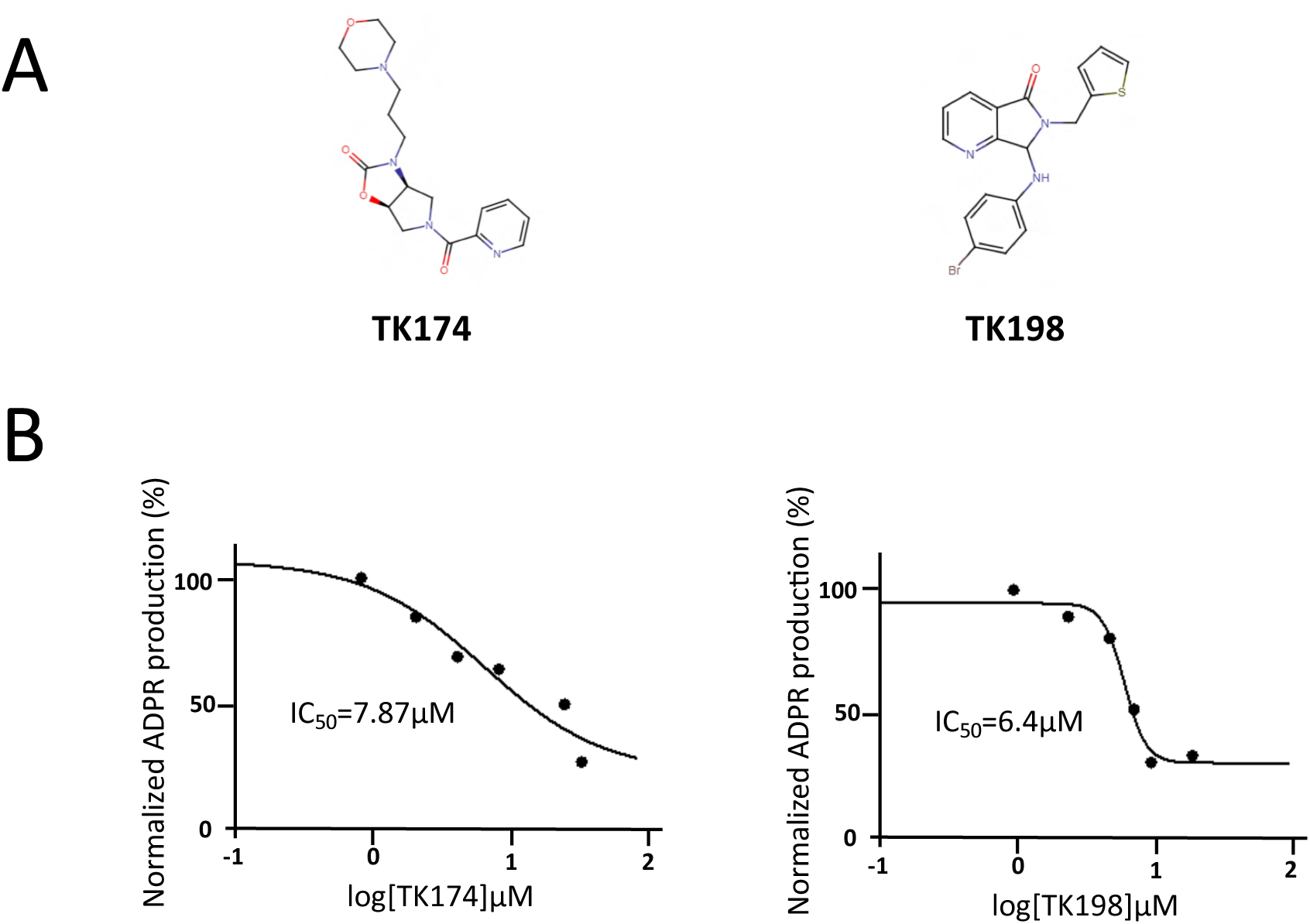
A) Chemical structure of 2 additional hSARM1 inhibitory compounds. B) Determination of IC_50_ values, as in figure 3.

**Table 1:**
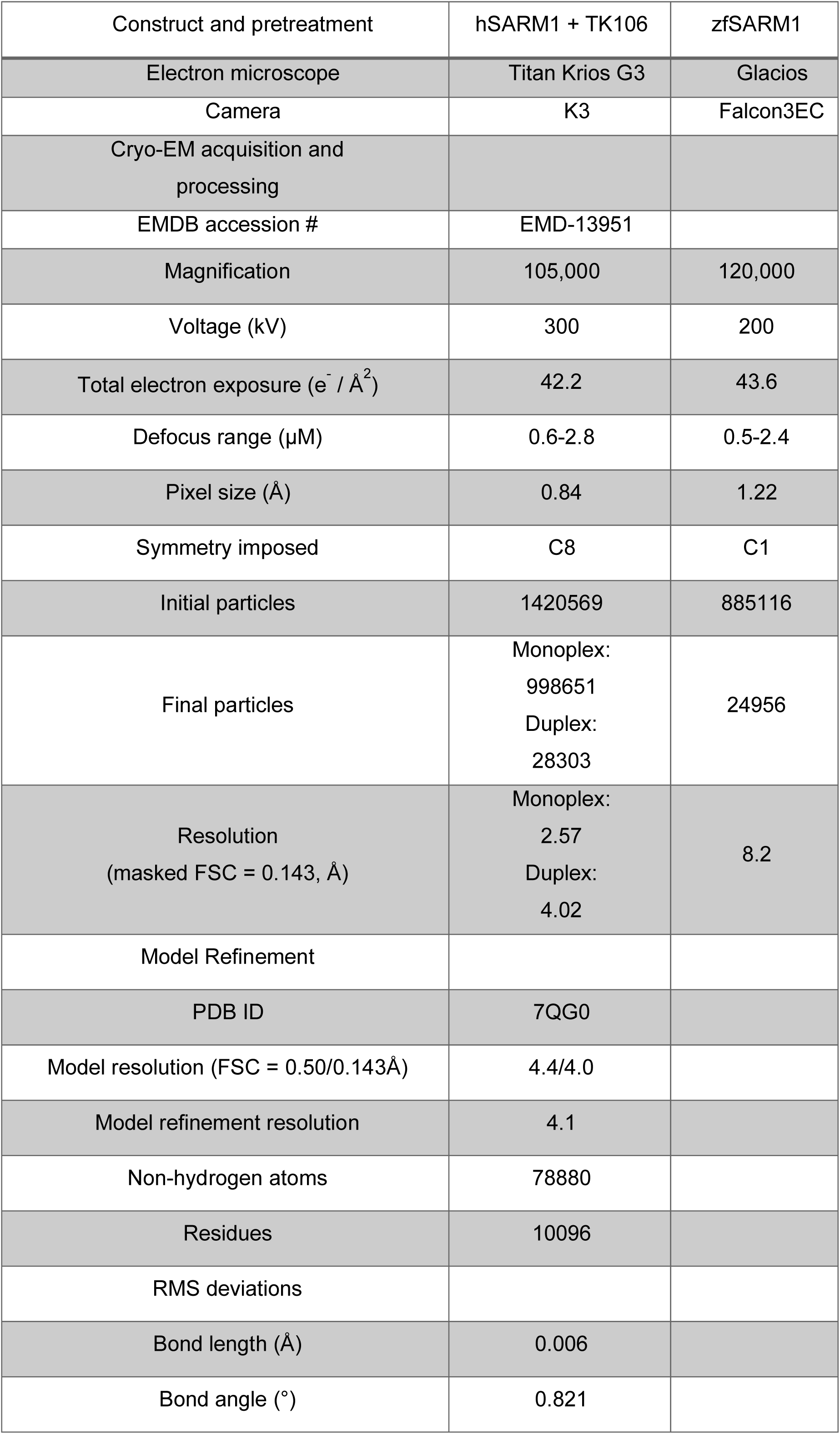
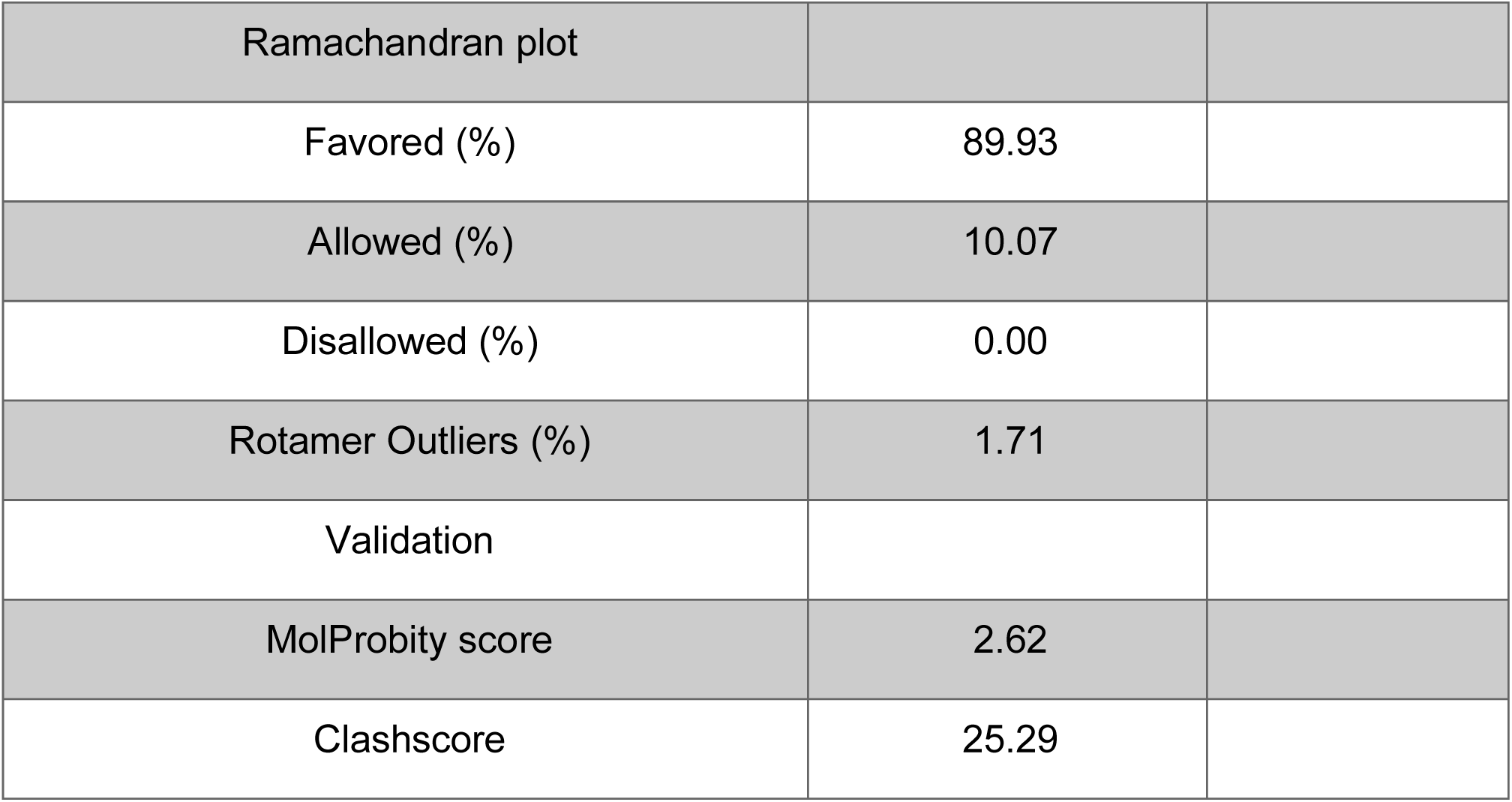
Cryo-EM data acquisition, reconstruction and model refinement statistics.

|  |  |  |
| --- | --- | --- |
| Construct and pretreatment | hSARM1 + TK106 | zfSARM1 |
| Electron microscope | Titan Krios G3 | Glacios |
| Camera | K3 | Falcon3EC |
| Cryo-EM acquisition and processing |  |  |
| EMDB accession # | EMD-13951 |  |
| Magnification | 105,000 | 120,000 |
| Voltage (kV) | 300 | 200 |
| Total electron exposure ($e^- / \text{\AA}^2$) | 42.2 | 43.6 |
| Defocus range ($\mu\text{M}$) | 0.6-2.8 | 0.5-2.4 |
| Pixel size ($\text{\AA}$) | 0.84 | 1.22 |
| Symmetry imposed | C8 | C1 |
| Initial particles | 1420569 | 885116 |
| Final particles | Monoplex:<br>998651<br>Duplex:<br>28303 | 24956 |
| Resolution<br>(masked FSC = 0.143, $\text{\AA}$) | Monoplex:<br>2.57<br>Duplex:<br>4.02 | 8.2 |
| Model Refinement |  |  |
| PDB ID | 7QG0 |  |
| Model resolution (FSC = 0.50/0.143$\text{\AA}$) | 4.4/4.0 | |
| Model refinement resolution | 4.1 |  |
| Non-hydrogen atoms | 78880 |  |
| Residues | 10096 |  |
| RMS deviations |  |  |
| Bond length ($\text{\AA}$) | 0.006 | |
| Bond angle ($^\circ$) | 0.821 | |

|  |  |
| --- | --- |
| Ramachandran plot |  |
| Favored (%) | 89.93 |
| Allowed (%) | 10.07 |
| Disallowed (%) | 0.00 |
| Rotamer Outliers (%) | 1.71 |
| Validation |  |
| MolProbity score | 2.62 |
| Clashscore | 25.29 |

**Figure 5 supplement 1.**
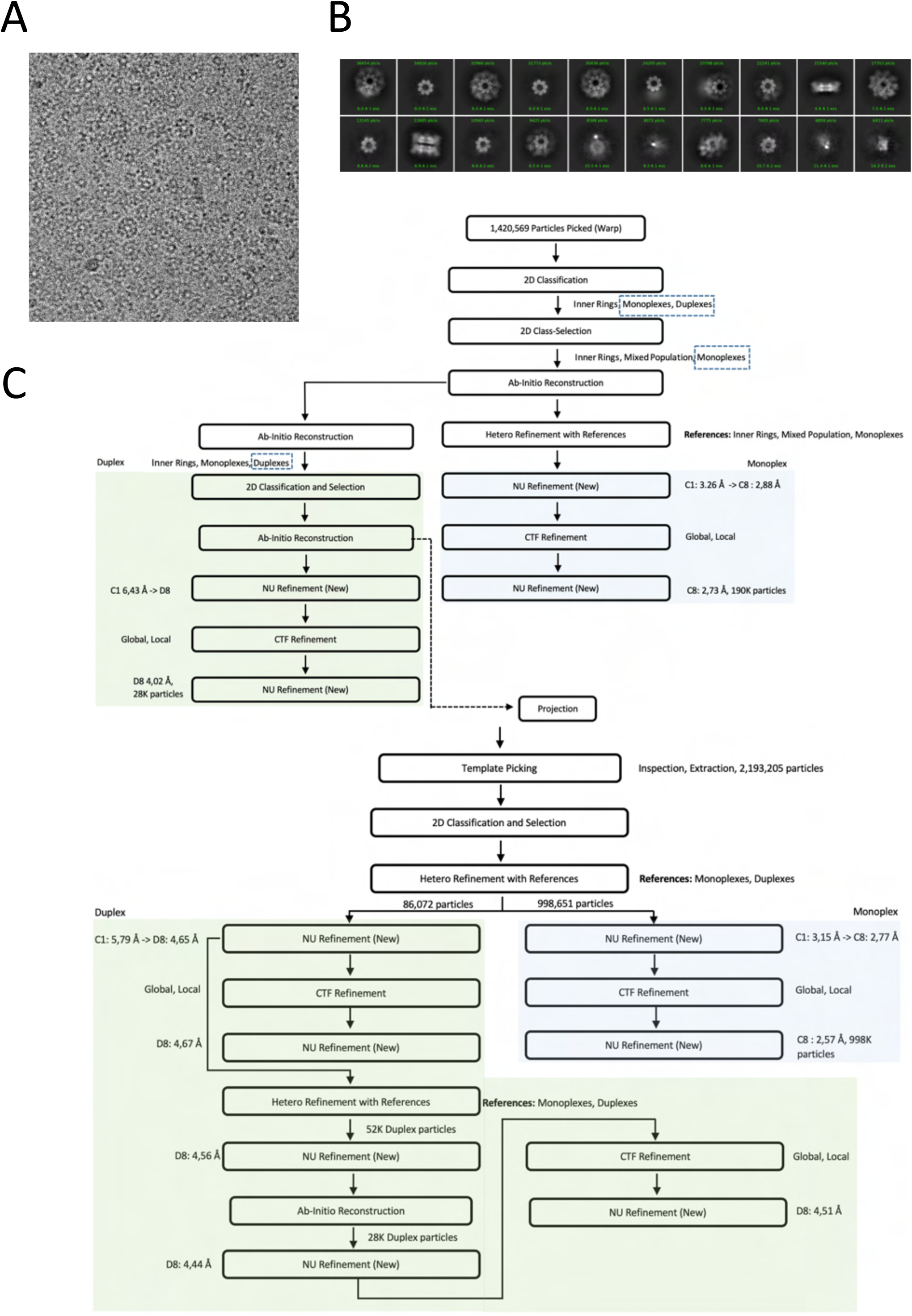
A) Representative cryo-EM micrograph of TK106 supplemented hSARM1. B) 2D class averages of the entire dataset. C) Flowchart of the cryo-EM processing steps of the entire dataset.

**Figure 5 supplement 2.**
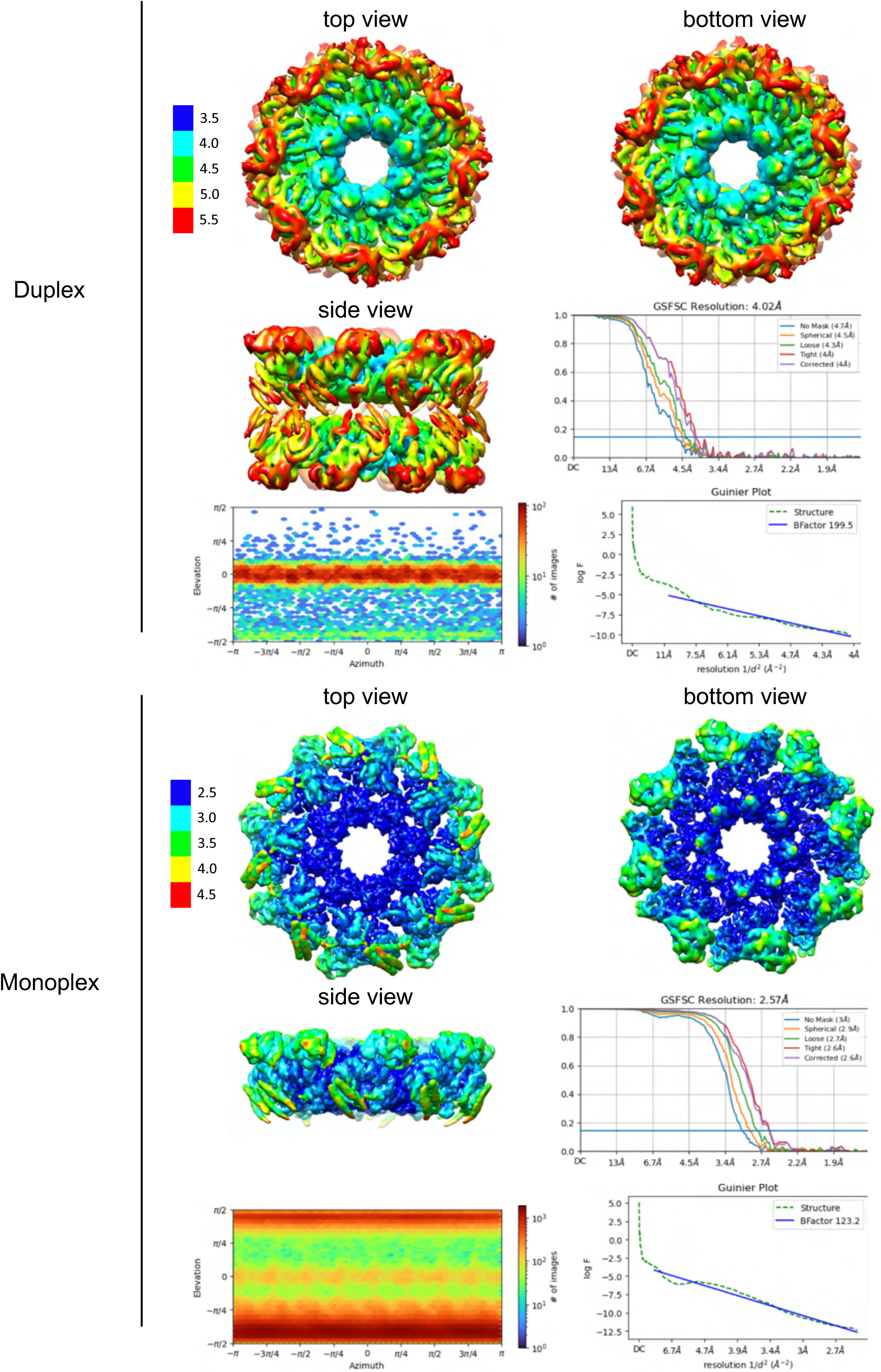
Resolution, angular distribution, and B-factor estimations of the cryo-EM maps of TK106 supplemented hSARM1 duplex and monoplex.

**Figure 5 supplement 3.**
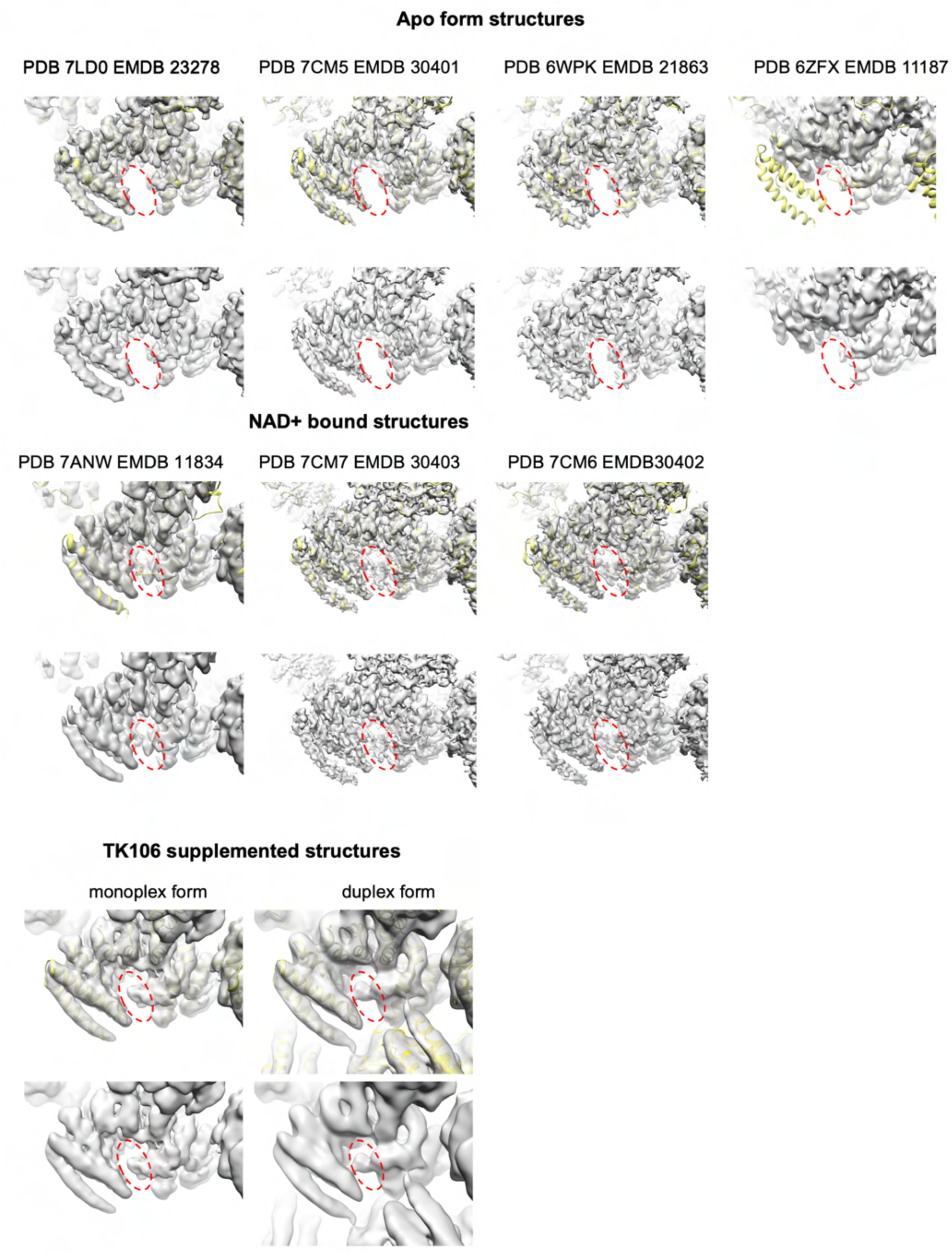
Comparison of available cryo-EM map densities of hSARM1, set to similar contour level at the ARM domain’s concave surface (Bratkowski et al., 2020; Figley et al., 2021; Jiang et al., 2020; Sporny et al., 2020). Dashed red oval marks the site of interest, highlighting the difference in map density between apo structures (empty site) and NAD+ or TK106 supplemented structures (occupied sites).

**Figure 6 supplement 1.**
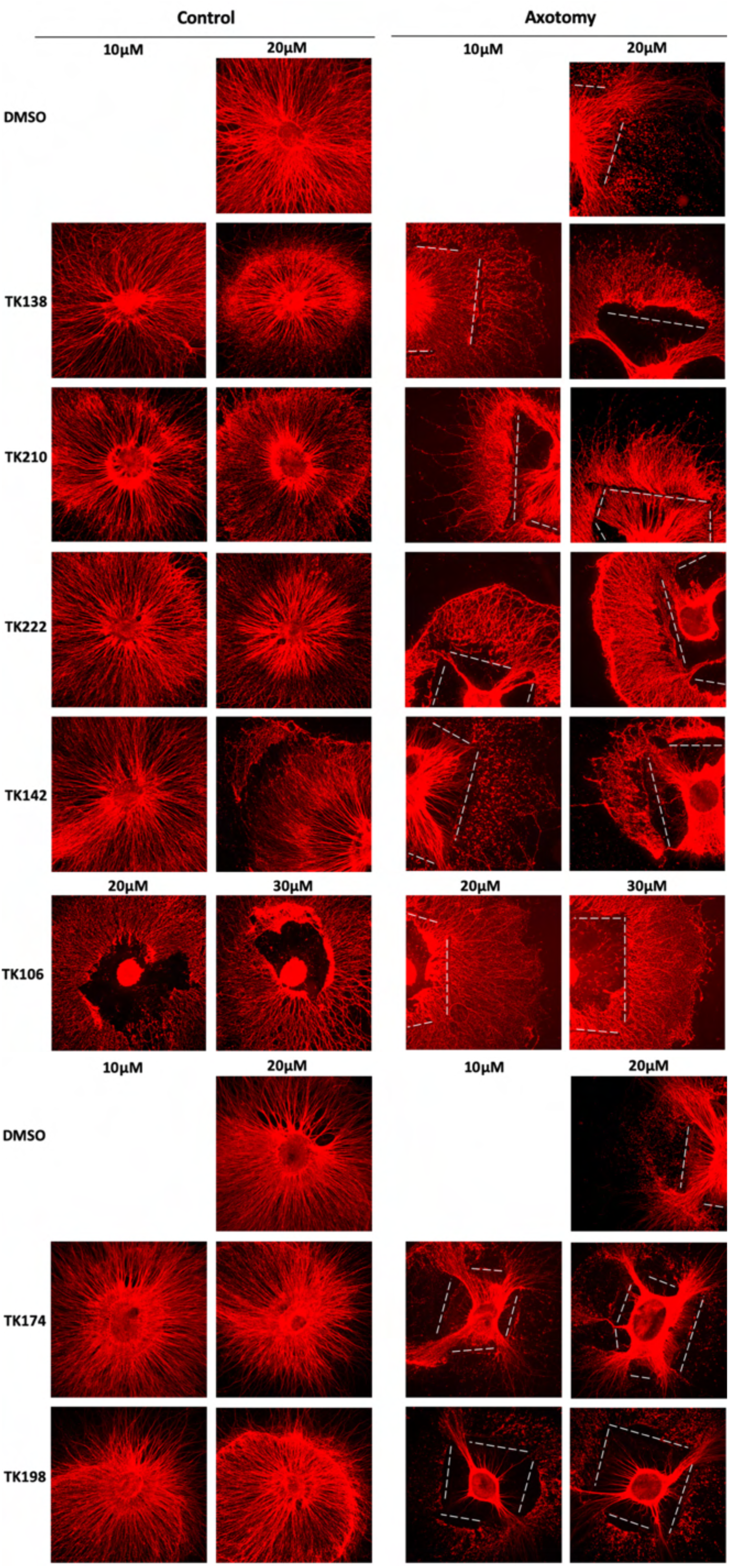
Extended representation of Fig. 6A. Protective and toxic effects of hSARM1 inhibitors over axon degeneration in mouse DRG explants.

## References

Afonine, P.V., Poon, B.K., Read, R.J., Sobolev, O.V., Terwilliger, T.C., Urzhumtsev, A., and Adams, P.D. (2018). Real-space refinement in PHENIX for cryo-EM and crystallography. Acta crystallographica Section D, Structural biology 74, 531–544.

Angeletti, C., Amici, A., Gilley, J., Loreto, A., Trapanotto, A.G., Antoniou, C., Coleman, M.P., and Orsomando, G. (2021). Programmed axon death executor SARM1 is a multi-functional NAD(P)ase with prominent base exchange activity, all regulated by physiological levels of NMN, NAD, NADP and other metabolites. bioRxiv.

Bosanac, T., Hughes, R.O., Engber, T., Devraj, R., Brearley, A., Danker, K., Young, K., Kopatz, J., Hermann, M., Berthemy, A., et al. (2021). Pharmacological SARM1 inhibition protects axon structure and function in paclitaxel-induced peripheral neuropathy. Brain.

Bratkowski, M., Xie, T., Thayer, D.A., Lad, S., Mathur, P., Yang, Y.S., Danko, G., Burdett, T.C., Danao, J., Cantor, A., et al. (2020). Structural and Mechanistic Regulation of the Pro-degenerative NAD Hydrolase SARM1. Cell reports 32, 107999.

Cambronne, X.A., Stewart, M.L., Kim, D., Jones-Brunette, A.M., Morgan, R.K., Farrens, D.L., Cohen, M.S., and Goodman, R.H. (2016). Biosensor reveals multiple sources for mitochondrial NAD(+). Science 352, 1474–1477.

Cetinkaya-Fisgin, A., Luan, X., Reed, N., Jeong, Y.E., Oh, B.C., and Hoke, A. (2020). Cisplatin induced neurotoxicity is mediated by Sarm1 and calpain activation. Sci Rep 10, 21889.

Chuang, C.F., and Bargmann, C.I. (2005). A Toll-interleukin 1 repeat protein at the synapse specifies asymmetric odorant receptor expression via ASK1 MAPKKK signaling. Genes Dev 19, 270–281.

Daina, A., Michielin, O., and Zoete, V. (2017). SwissADME: a free web tool to evaluate pharmacokinetics, drug-likeness and medicinal chemistry friendliness of small molecules. Sci Rep 7, 42717.

Emsley, P., Lohkamp, B., Scott, W.G., and Cowtan, K. (2010). Features and development of Coot. Acta Cryst D 66, 486–501.

Essuman, K., Summers, D.W., Sasaki, Y., Mao, X., DiAntonio, A., and Milbrandt, J. (2017). The SARM1 Toll/Interleukin-1 Receptor Domain Possesses Intrinsic NAD(+) Cleavage Activity that Promotes Pathological Axonal Degeneration. Neuron 93, 1334–1343 e1335.

Figley, M.D., Gu, W., Nanson, J.D., Shi, Y., Sasaki, Y., Cunnea, K., Malde, A.K., Jia, X., Luo, Z., Saikot, F.K., et al. (2021). SARM1 is a metabolic sensor activated by an increased NMN/NAD(+) ratio to trigger axon degeneration. Neuron 109, 1118–1136 e1111.

Geisler, S., Doan, R.A., Strickland, A., Huang, X., Milbrandt, J., and DiAntonio, A. (2016). Prevention of vincristine-induced peripheral neuropathy by genetic deletion of SARM1 in mice. Brain 139, 3092–3108.

Gerdts, J., Brace, E.J., Sasaki, Y., DiAntonio, A., and Milbrandt, J. (2015). SARM1 activation triggers axon degeneration locally via NAD(+) destruction. Science 348, 453–457.

Gerdts, J., Summers, D.W., Milbrandt, J., and DiAntonio, A. (2016). Axon Self-Destruction: New Links among SARM1, MAPKs, and NAD+ Metabolism. Neuron 89, 449–460.

Gerdts, J., Summers, D.W., Sasaki, Y., DiAntonio, A., and Milbrandt, J. (2013). Sarm1-mediated axon degeneration requires both SAM and TIR interactions. J Neurosci 33, 13569–13580.

Gilley, J., Jackson, O., Pipis, M., Estiar, M.A., Al-Chalabi, A., Danzi, M.C., van Eijk, K.R., Goutman, S.A., Harms, M.B., Houlden, H., et al. (2021). Enrichment of SARM1 alleles encoding variants with constitutively hyperactive NADase in patients with ALS and other motor nerve disorders. Elife 10.

Guez-Haddad, J., Sporny, M., Sasson, Y., Gevorkyan-Airapetov, L., Lahav-Mankovski, N., Margulies, D., Radzimanowski, J., and Opatowsky, Y. (2015). The Neuronal Migration Factor srGAP2 Achieves Specificity in Ligand Binding through a Two-Component Molecular Mechanism. Structure 23, 1989–2000.

Hara, N., Osago, H., Hiyoshi, M., Kobayashi-Miura, M., and Tsuchiya, M. (2019). Quantitative analysis of the effects of nicotinamide phosphoribosyltransferase induction on the rates of NAD+ synthesis and breakdown in mammalian cells using stable isotope-labeling combined with mass spectrometry. PLoS One 14, e0214000.

Horsefield, S., Burdett, H., Zhang, X., Manik, M.K., Shi, Y., Chen, J., Qi, T., Gilley, J., Lai, J.S., Rank, M.X., et al. (2019). NAD(+) cleavage activity by animal and plant TIR domains in cell death pathways. Science 365, 793–799.

Houtkooper, R.H., Canto, C., Wanders, R.J., and Auwerx, J. (2010). The secret life of NAD+: an old metabolite controlling new metabolic signaling pathways. Endocr Rev 31, 194–223.

Hughes, R.O., Bosanac, T., Mao, X., Engber, T.M., DiAntonio, A., Milbrandt, J., Devraj, R., and Krauss, R. (2021). Small Molecule SARM1 Inhibitors Recapitulate the SARM1(-/-) Phenotype and Allow Recovery of a Metastable Pool of Axons Fated to Degenerate. Cell reports 34, 108588.

Jiang, Y., Liu, T., Lee, C.H., Chang, Q., Yang, J., and Zhang, Z. (2020). The NAD(+)-mediated self-inhibition mechanism of pro-neurodegenerative Sarm1. Nature.

Kandiah, E., Giraud, T., de Maria Antolinos, A., Dobias, F., Effantin, G., Flot, D., Hons, M., Schoehn, G., Susini, J., Svensson, O., et al. (2019). CM01: a facility for cryo-electron microscopy at the European Synchrotron. Acta crystallographica Section D, Structural biology 75, 528–535.

Kim, Y., Zhou, P., Qian, L., Chuang, J.Z., Lee, J., Li, C., Iadecola, C., Nathan, C., and Ding, A. (2007). MyD88-5 links mitochondria, microtubules, and JNK3 in neurons and regulates neuronal survival. J Exp Med 204, 2063–2074.

Ko, K.W., Milbrandt, J., and DiAntonio, A. (2020). SARM1 acts downstream of neuroinflammatory and necroptotic signaling to induce axon degeneration. J Cell Biol 219.

Krissinel, E. (2012). Enhanced fold recognition using efficient short fragment clustering. J Mol Biochem 1, 76–85.

Langer, G., Cohen, S.X., Lamzin, V.S., and Perrakis, A. (2008). Automated macromolecular model building for X-ray crystallography using ARP/wARP version 7. Nature protocols 3, 1171–1179.

Lebedev, A.A., Young, P., Isupov, M.N., Moroz, O.V., Vagin, A.A., and Murshudov, G.N. (2012). JLigand: a graphical tool for the CCP4 template-restraint library. Acta Crystallogr D Biol Crystallogr 68, 431–440.

Li, W.H., Huang, K., Cai, Y., Wang, Q.W., Zhu, W.J., Hou, Y.N., Wang, S., Cao, S., Zhao, Z.Y., Xie, X.J., et al. (2021). Permeant fluorescent probes visualize the activation of SARM1 and uncover an anti-neurodegenerative drug candidate. Elife 10.

Li, X., Mooney, P., Zheng, S., Booth, C.R., Braunfeld, M.B., Gubbens, S., Agard, D.A., and Cheng, Y. (2013). Electron counting and beam-induced motion correction enable near-atomic-resolution single-particle cryo-EM. Nature methods 10, 584–590.

Liebschner, D., Afonine, P.V., Baker, M.L., Bunkoczi, G., Chen, V.B., Croll, T.I., Hintze, B., Hung, L.W., Jain, S., McCoy, A.J., et al. (2019). Macromolecular structure determination using X-rays, neutrons and electrons: recent developments in Phenix. Acta crystallographica Section D, Structural biology 75, 861–877.

Liu, H.W., Smith, C.B., Schmidt, M.S., Cambronne, X.A., Cohen, M.S., Migaud, M.E., Brenner, C., and Goodman, R.H. (2018a). Pharmacological bypass of NAD(+) salvage pathway protects neurons from chemotherapy-induced degeneration. Proc Natl Acad Sci U S A 115, 10654–10659.

Liu, L., Su, X., Quinn, W.J., 3rd, Hui, S., Krukenberg, K., Frederick, D.W., Redpath, P., Zhan, L., Chellappa, K., White, E., et al. (2018b). Quantitative Analysis of NAD Synthesis-Breakdown Fluxes. Cell Metab 27, 1067–1080 e1065.

Loreto, A., Angeletti, C., Gilley, J., Arthur-Farraj, P., Merlini, E., Amici, A., Desrochers, L.M., Wang, Q., Orsomando, G., and Coleman, M.P. (2020). Potent activation of SARM1 by NMN analogue VMN underlies vacor neurotoxicity. Preprint.

Loring, H.S., Parelkar, S.S., Mondal, S., and Thompson, P.R. (2020). Identification of the first noncompetitive SARM1 inhibitors. Bioorg Med Chem 28, 115644.

Ma, S., Lapin, D., Liu, L., Sun, Y., Song, W., Zhang, X., Logemann, E., Yu, D., Wang, J., Jirschitzka, J., et al. (2020). Direct pathogen-induced assembly of an NLR immune receptor complex to form a holoenzyme. Science 370.

Martin, R., Qi, T., Zhang, H., Liu, F., King, M., Toth, C., Nogales, E., and Staskawicz, B.J. (2020). Structure of the activated ROQ1 resistosome directly recognizing the pathogen effector XopQ. Science 370.

Martinez, M., Jimenez-Moreno, A., Maluenda, D., Ramirez-Aportela, E., Melero, R., Cuervo, A., Conesa, P., Del Cano, L., Fonseca, Y.C., Sanchez-Garcia, R., et al. (2020). Integration of Cryo-EM Model Building Software in Scipion. J Chem Inf Model.

Murshudov, G.N., Skubak, P., Lebedev, A.A., Pannu, N.S., Steiner, R.A., Nicholls, R.A., Winn, M.D., Long, F., and Vagin, A.A. (2011). REFMAC5 for the refinement of macromolecular crystal structures. Acta Cryst D 67, 355–367.

Osterloh, J.M., Yang, J., Rooney, T.M., Fox, A.N., Adalbert, R., Powell, E.H., Sheehan, A.E., Avery, M.A., Hackett, R., Logan, M.A., et al. (2012). dSarm/Sarm1 is required for activation of an injury-induced axon death pathway. Science 337, 481–484.

Ozaki, E., Gibbons, L., Neto, N.G., Kenna, P., Carty, M., Humphries, M., Humphries, P., Campbell, M., Monaghan, M., Bowie, A., et al. (2020). SARM1 deficiency promotes rod and cone photoreceptor cell survival in a model of retinal degeneration. Life Sci Alliance 3.

Punjani, A., Rubinstein, J.L., Fleet, D.J., and Brubaker, M.A. (2017). cryoSPARC: algorithms for rapid unsupervised cryo-EM structure determination. Nature Methods 14, 290–+.

Sasaki, Y., Kakita, H., Kubota, S., Sene, A., Lee, T.J., Ban, N., Dong, Z., Lin, J.B., Boye, S.L., DiAntonio, A., et al. (2020). SARM1 depletion rescues NMNAT1-dependent photoreceptor cell death and retinal degeneration. Elife 9.

Sasaki, Y., Zhu, J., Shi, Y., Gu, W., Kobe, B., Ve, T., DiAntonio, A., and Milbrandt, J. (2021). Nicotinic acid mononucleotide is an allosteric SARM1 inhibitor promoting axonal protection. Exp Neurol 345, 113842.

Shacham-Silverberg, V., Sar Shalom, H., Goldner, R., Golan-Vaishenker, Y., Gurwicz, N., Gokhman, I., and Yaron, A. (2018). Phosphatidylserine is a marker for axonal debris engulfment but its exposure can be decoupled from degeneration. Cell Death Dis 9, 1116.

Shen, C., Vohra, M., Zhang, P., Mao, X., Figley, M.D., Zhu, J., Sasaki, Y., Wu, H., DiAntonio, A., and Milbrandt, J. (2021). Multiple domain interfaces mediate SARM1 autoinhibition. Proc Natl Acad Sci U S A 118.

Sporny, M., Guez-Haddad, J., Khazma, T., Yaron, A., Dessau, M., Shkolnisky, Y., Mim, C., Isupov, M.N., Zalk, R., Hons, M., et al. (2020). Structural basis for SARM1 inhibition and activation under energetic stress. Elife 9.

Sporny, M., Guez-Haddad, J., Lebendiker, M., Ulisse, V., Volf, A., Mim, C., Isupov, M.N., and Opatowsky, Y. (2019). Structural Evidence for an Octameric Ring Arrangement of SARM1. J Mol Biol 431, 3591–3605.

Summers, D.W., Gibson, D.A., DiAntonio, A., and Milbrandt, J. (2016). SARM1-specific motifs in the TIR domain enable NAD+ loss and regulate injury-induced SARM1 activation. Proc Natl Acad Sci U S A 113, E6271–E6280.

Tal, N., Morehouse, B.R., Millman, A., Stokar-Avihail, A., Avraham, C., Fedorenko, T., Yirmiya, E., Herbst, E., Brandis, A., Mehlman, T., et al. (2021). Cyclic CMP and cyclic UMP mediate bacterial immunity against phages. Cell.

Tegunov, D., and Cramer, P. (2019). Real-time cryo-electron microscopy data preprocessing with Warp. Nat Methods 16, 1146–1152.

Tong, L. (2021). How to diSARM the executioner of axon degeneration. Nat Struct Mol Biol 28, 10–12.

Turkiew, E., Falconer, D., Reed, N., and Hoke, A. (2017). Deletion of Sarm1 gene is neuroprotective in two models of peripheral neuropathy. J Peripher Nerv Syst 22, 162–171.

Uccellini, M.B., Bardina, S.V., Sanchez-Aparicio, M.T., White, K.M., Hou, Y.J., Lim, J.K., and Garcia-Sastre, A. (2020). Passenger Mutations Confound Phenotypes of SARM1-Deficient Mice. Cell reports 31, 107498.

Vagin, A., and Teplyakov, A. (2010). Molecular replacement with MOLREP. Acta Cryst D 66, 22–25.

Winn, M.D., Ballard, C.C., Cowtan, K.D., Dodson, E.J., Emsley, P., Evans, P.R., Keegan, R.M., Krissinel, E.B., Leslie, A.G., McCoy, A., et al. (2011). Overview of the CCP4 suite and current developments. Acta Cryst D 67, 235–242.

Wu, T., Zhu, J., Strickland, A., Ko, K.W., Sasaki, Y., Dingwall, C., Yamada, Y., Figley, M.D., Mao, X., Neiner, A., et al. (2021). Neurotoxins subvert the allosteric activation mechanism of SARM1 to induce neuronal loss. preprint.

Yom-Tov, G., Barak, R., Matalon, O., Barda-Saad, M., Guez-Haddad, J., and Opatowsky, Y. (2017). Robo Ig4 Is a Dimerization Domain. J Mol Biol 429, 3606–3616.

Zhang, K. (2016). Gctf: Real-time CTF determination and correction. J Struct Biol 193, 1–12.

Zhao, Z.Y., Xie, X.J., Li, W.H., Liu, J., Chen, Z., Zhang, B., Li, T., Li, S.L., Lu, J.G., Zhang, L., et al. (2019). A Cell-Permeant Mimetic of NMN Activates SARM1 to Produce Cyclic ADP-Ribose and Induce Non-apoptotic Cell Death. iScience 15, 452–466.

